# Activation of automethylated PRC2 by dimerization on chromatin

**DOI:** 10.1101/2023.10.12.562141

**Authors:** Paul V. Sauer, Egor Pavlenko, Trinity Cookis, Linda C. Zirden, Juliane Renn, Ankush Singhal, Pascal Hunold, Michaela N. Hoehne, Olivia van Ray, Robert Hänsel-Hertsch, Karissa Y. Sanbonmatsu, Eva Nogales, Simon Poepsel

## Abstract

Polycomb Repressive Complex 2 (PRC2) is an epigenetic regulator that trimethylates lysine 27 of histone 3 (H3K27me3) and is essential for embryonic development and cellular differentiation. H3K27me3 is associated with transcriptionally repressed chromatin and is established when PRC2 is allosterically activated upon methyl-lysine binding by the regulatory subunit EED. Automethylation of the catalytic subunit EZH2 stimulates its activity by an unknown mechanism. Here, we show that PRC2 forms a dimer on chromatin in which an inactive, automethylated PRC2 protomer is the allosteric activator of a second PRC2 that is poised to methylate H3 of a substrate nucleosome. Functional assays support our model of allosteric *trans*-autoactivation via EED, suggesting a novel mechanism mediating context-dependent activation of PRC2. Our work showcases the molecular mechanism of auto-modification coupled dimerization in the regulation of chromatin modifying complexes.

## Introduction

Polycomb Repressive Complex 2 (PRC2) is an essential chromatin regulator that harbors histone methyltransferase (HMTase) activity mediated by its catalytic subunit Enhancer of Zeste Homolog 2 (EZH2). PRC2 is the only known factor that establishes the trimethylation of lysine 27 of histone H3 (H3K27me3), a chromatin mark associated with transcriptional repression and whose tight spatiotemporal regulation is essential during development and differentiation. In addition to EZH2 (or its less active homolog EZH1), the core PRC2 complex consists of SUZ12, RBAP46/48 and EED.^1^ The binding of EED to trimethylated-lysine bearing peptides is central to PRC2 activity, since it allosterically activates EZH2, a requirement for efficient trimethylation of H3K27.^2^ Upon engagement of specific methylated peptides by EED, a flexible loop of EZH2, the stimulatory-response motif (SRM), folds into a short α-helix and stabilizes an active conformation of EZH2 by bridging the EED ligand binding site and the critical SET-I helix of the EZH2 SET domain.^3,4^ Two EED-binding ligands, themselves trimethylated by PRC2, have been described as functional activators: JARID2, an accessory PRC2 subunit that contributes to the targeting of the complex and *de novo* deposition of H3K27me3, and that binds EED through trimethylated JARID2 K116^5,6^; and H3K27me3 itself, which facilitates H3K27me3 propagation in a positive feedback loop. Highlighting the importance of this regulatory, allosteric mechanism, H3K27me3 and gene expression dynamics are impaired during development when EED-EZH2 communication is interrupted.^2,7,8^

Just like kinase activation can occur upon autophosphorylation,^9,10^ chromatin-modifying enzymes can also act upon themselves for regulation,^11,12^ sometimes in a manner that is linked to a change in oligomeric state (e.g. transcription factor-mediated dimerization and auto-acetylation of P300 has been suggested to be critical for target specificity and to act as molecular short-term memory during inflammatory response^13^). Recently, EZH2 has been shown to be automethylated in *cis*, leading to increased HMTase activity through a yet unknown mechanism.^14,15^ The three automethylated EZH2 residues, K510, K514 and K515, are part of a flexible loop (hereafter the automethylation loop, “am-loop”) that fold into an α-helix upon engagement with nucleosome substrates.^8,16^ This region is referred to as the “bridge helix” because it bridges the nucleosomal DNA, the substrate histone H3 tail, and the EZH2 SET domain.^8,16^

Here, we set out to investigate how automethylation of EZH2 may activate human PRC2 using single-particle cryo-electron microscopy (cryo-EM) and functional assays. To this end, we studied the structural implications of substrate chromatin engagement and methylation in the absence of any allosteric activator other than automethylated EZH2, i.e. using mono- nucleosomes and recombinant PRC2 lacking the stimulatory subunit JARID2. We show that in a novel chromatin- and automethylation-dependent manner, PRC2 dimerizes such that an automethylated inactive PRC2 complex serves as an allosteric activator via the EED regulatory site of a second substrate-engaged PRC2. Using separation of function mutants, we provide evidence that automethylation indeed regulates PRC2 activity in *trans* and that it functions in defined genomic contexts. Taken together, we propose that dimerization-dependent stimulation of PRC2 HMTase activity in *trans*, driven by EZH2 automethylation and local PRC2 concentration, represents a new mechanism to regulate the establishment of H3K27me3 heterochromatin domains.

## Results

### Dimerization of automethylated PRC2 on chromatin

To study the mechanism of EZH2 automethylation mediated activation, we set out to solve the cryo-EM structure of PRC2 engaged with a substrate nucleosome in the absence of H3K27me3 or JARID2, and therefore with the am-loop as its only source of methylated peptides for possible allosteric activation. Mass spectrometry analysis of recombinantly expressed human PRC2, composed of EZH2, SUZ12, EED, RBAP48 and AEBP2 revealed EZH2 K510, K514 and K515 to be methylated to varying degrees. Incubation of recombinant PRC2 with S-adenosyl methionine (SAM) cofactor further increased the levels of am-loop automethylation (Figure S1). Cryo-EM analysis of automethylated PRC2 incubated with nucleosomes containing 40 bp stretches of linker DNA on both sides resolved two distinct species: first, nucleosomes bound by a single PRC2 complex (Figure S2), and second, two PRC2 complexes interacting with one nucleosome and with each other in a non-symmetrical dimer (Figure 1, Figure S3). Single nucleosome engaged PRC2 complexes were in a basal state in which PRC2 is not allosterically activated, as we previously described^16^ (i.e. folded bridge helix, but unfolded SRM, see later and Figure 4). Here, we will focus mainly on the observed PRC2 dimer-nucleosome complex (Figure 1A-B, Figure S2-5, Figure S3, Table S1).

**Figure 1.**
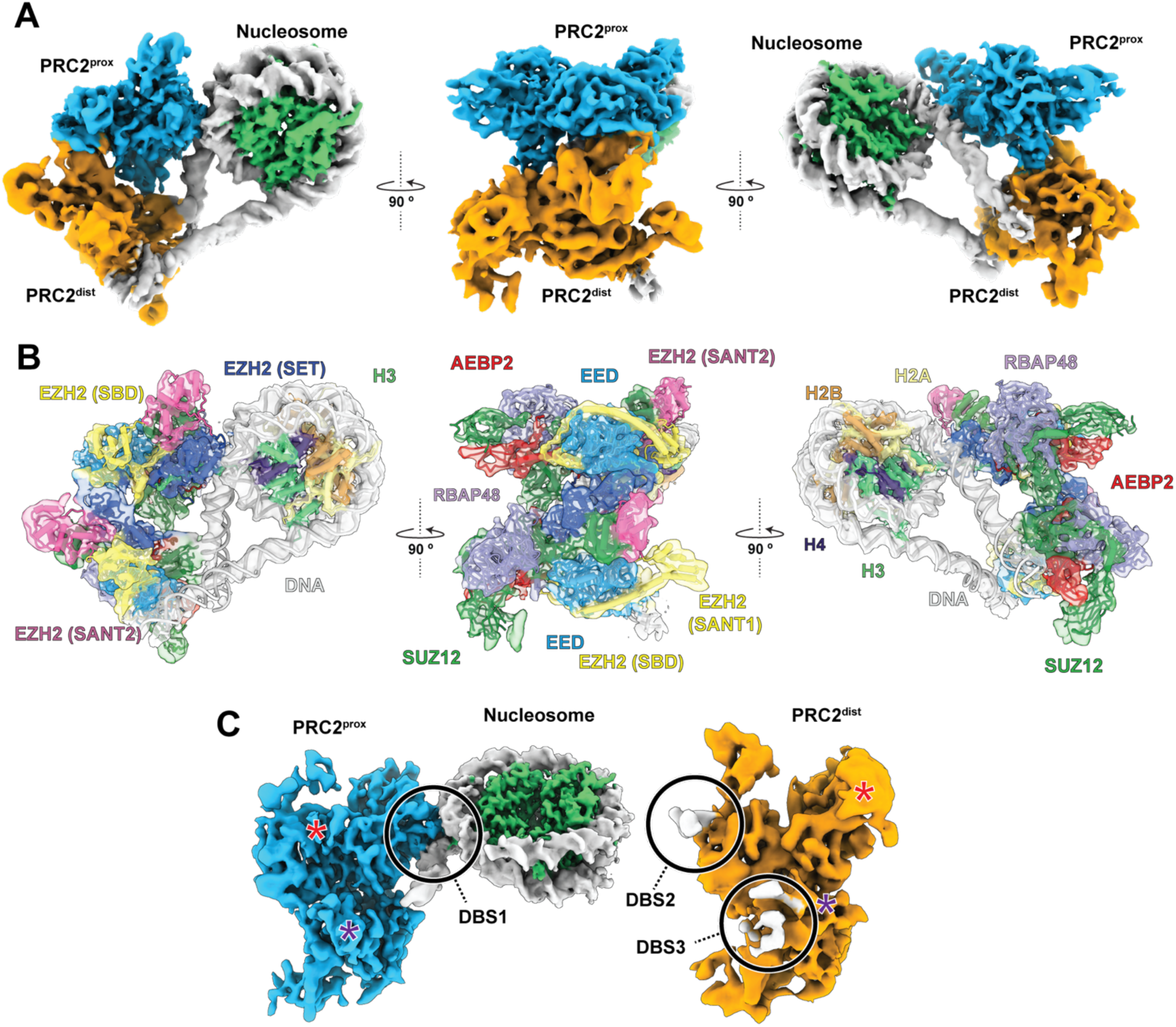
Characterization of an asymmetric PRC2 dimer bound to a nucleosome. (A) Composite cryo-EM density for proximal (blue) and distal (orange) PRC2 complexes simultaneously engaged with a nucleosome shown from three orthogonal orientations. (B) As A, but with docked models colored by PRC2 subunit/domain (same colors used throughout). (C) The structure in A has been rotated and “opened” to show the proximal and distal PRC2 separated and in the same general orientation. DNA binding sites (DBS) are indicated (see also Figure 2). Asterisks mark the dimer interfaces between EED^prox^ and EZH2^dist^ (red) and between SUZ12^prox^ and SUZ12^dist^ (purple).

The two PRC2 complexes and the nucleosome form a flexible, tripartite structure stabilized by contacts between the three components (Figure 1). In contrast to other recently described symmetric PRC2 dimers,^17,18^ the two PRC2 complexes are arranged asymmetrically, with only one of them contacting the histone octamer in a way that is compatible with H3 methylation. We refer to this PRC2 as nucleosome-proximal PRC2 (PRC2^prox^) (Figure 1A,C, blue). The second PRC2 is located distal from the histone core (referred to as PRC2^dist^) (Figure 1A,C, orange) and interacts distinctly with each of the two DNA linkers, as well as with the PRC2^prox^, but not with the nucleosome core (Figure 1). Employing 3D Flexible refinement (3DFlex),^19^ we were able to characterize the extensive motions within this molecular arrangement (Figure S4). Despite the flexibility and transient nature of the complex, this strategy enabled us to unambiguously fit PRC2 and nucleosome models into an improved density map (Figure 2B), defining the inter- PRC2 and PRC2-DNA interfaces, as well as the distinct conformational states of the PRC2^prox^ and PRC2^dist^.

**Figure 2.**
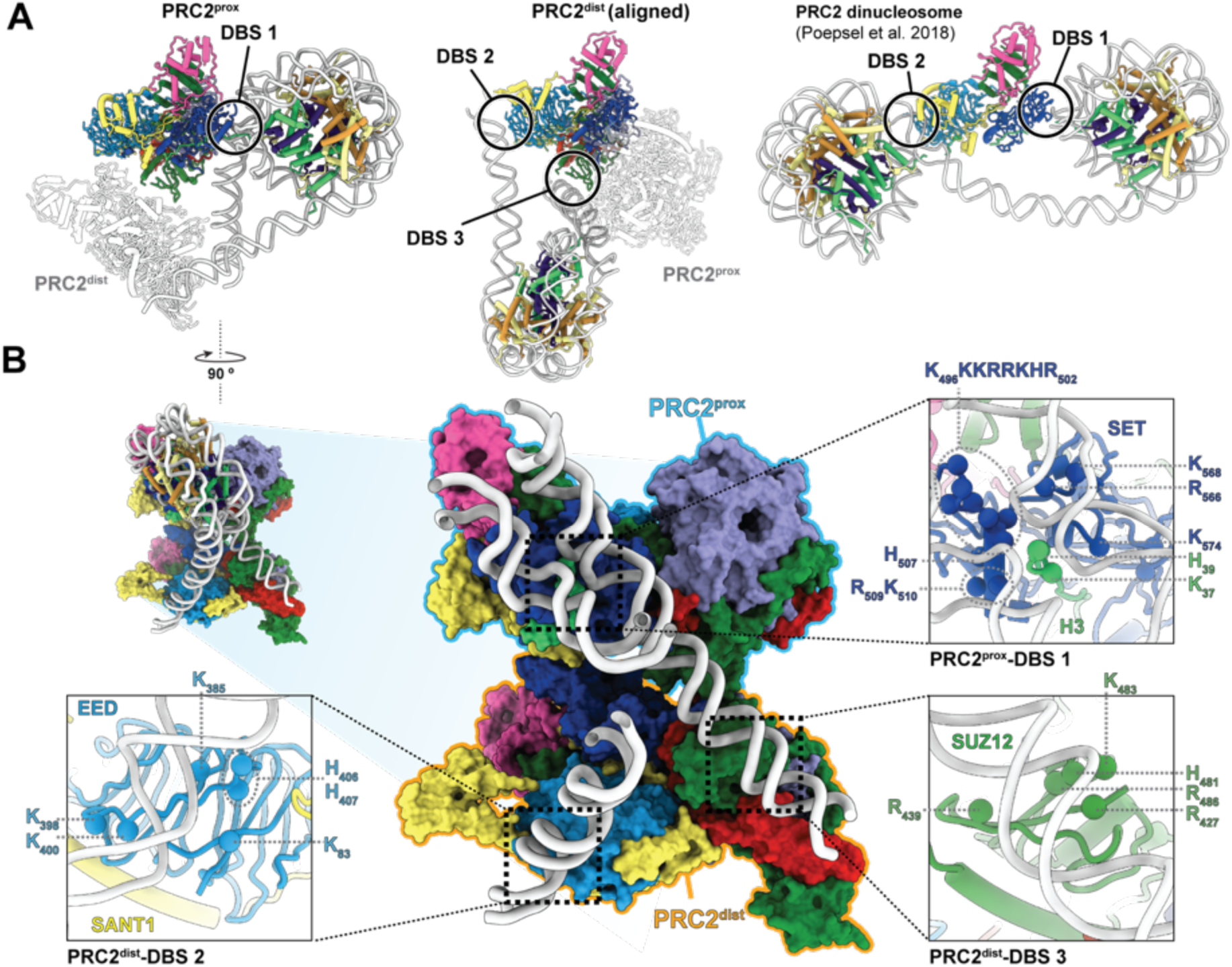
Three distinct DNA binding sites are present in the PRC2 dimer. (A) DNA interaction sites for PRC2^prox^ (left), PRC2^dist^ (center) and for PRC2 engaged with a dinucleosome (from ^8^) (right) with all PRC2 complexes aligned to one another. The three distinct DNA binding sites (DBS) collectively observed are indicated as DBS1, DBS2, and DBS3. (B) PRC2 dimer viewed through the nucleosome (center; top left shows the relative orientation of the view shown with respect to the left panel in (A) before removing the histones and part of the DNA). The three different DBSs are displayed in more detail in the three zoom-out boxes. Positively charged residues within 10 Å of nucleic acid have been marked with spheres as possible sites of interaction with the phosphate backbone of the DNA.

In the PRC2^prox^, the SET domain of EZH2 engages the substrate H3 tail while its CXC domain contacts the nucleosomal DNA, as previously seen.^8,17,20^ We refer to this DNA binding site (DBS), which involves the bridge helix containing the automethylated lysines, as DBS1 (Figure 2A, left; Figure 2B). In the PRC2^dist^, a lateral surface of EED^dist^, which we refer to as DNA binding site 2 (DBS2), binds one DNA linker (Figure 2A, center; Figure 2B). The same site has been seen to bind DNA in the context of a hetero-dinucleosome substrate in which DBS2 binding of a H3K27me3 bearing nucleosome stimulates methylation of a substrate nucleosome engaged via DBS1 (Figure 2A, right).^8^ Thus, DBS2 in EED can engage either nucleosomal or linker DNA depending on the context. We additionally observe an unassigned cylindrical density contacted by a positively charged surface corresponding approximately to DBS2 in PRC2^prox^ (Figure S5) that may correspond to double stranded DNA and is absent in the PRC2 monomer reconstruction (Figure S5B). PRC2^dist^ also contacts the other linker DNA, in this case via the neck region of SUZ12^dist^, a novel DNA binding site of PRC2 that we refer to as DBS3 (Figure 2A, center; Figure 2B). Based on our docking into the cryo-EM density of PRC2^dist^, this DNA-binding interface likely involves the loop H_481_PKGA_485_ and several nearby positively charged residues of SUZ12 (Figure 2B). Therefore, PRC2 can utilize three distinct DNA binding surfaces (DBS1-3) to mediate interactions with the local chromatin environment, and the three of them are involved in the tripartite engagement described in this study.

### Interaction between PRC2^prox^ and PRC2^dist^ and allosteric activation via the am-loop

In addition to their interactions with the nucleosome, the PRC2^dist^ and PRC2^prox^ interact with each other via two different interfaces. A SUZ12-SUZ12 interface comprises loop H_481_PKGA_485_ and a stretch that includes L426 and R427 of SUZ12^prox^. These elements engage with a short, negatively charged helix in SUZ12^dist^ (E_542_FLESED_548_) (Figure 3A). Thus, the DBS3 of SUZ12 involving loop H_481_PKGA_485_ and nearby residues appear to mediate two different interactions important within the supramolecular arrangement we are describing: one in PRC2^dist^ with linker DNA, and one in PRC2^prox^ with another SUZ12 in PRC2^dist^.

**Figure 3.**
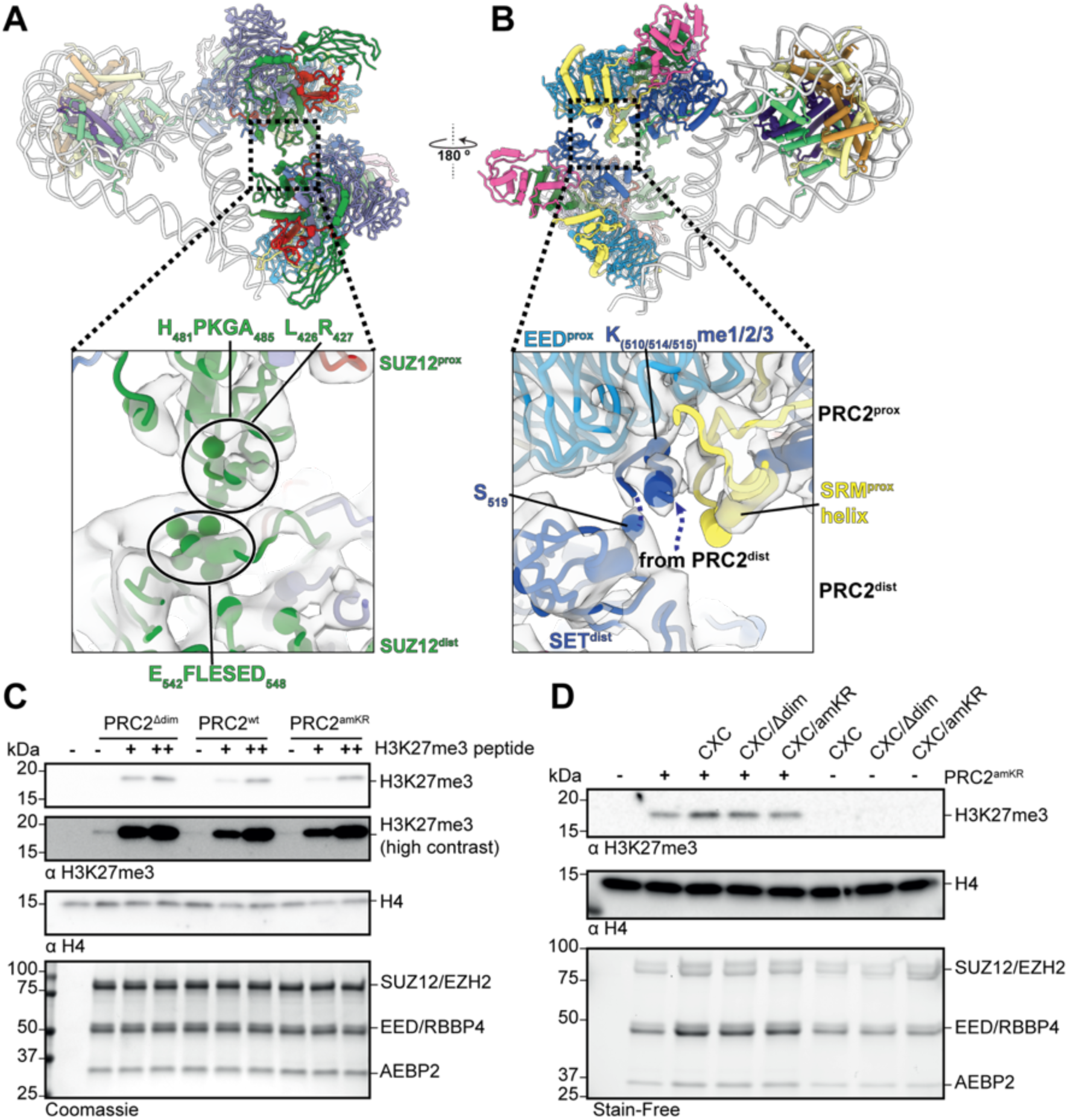
PRC2 dimer interfaces. (A) Dimer interface involving SUZ12^prox^ and SUZ12^dist^. (B) Interaction between the SET domain of the PRC2^dist^ with the EED of PRC2^prox^. Potential interacting residues based on proximity are indicated by spheres in the zoom-out panels. The dashed lines in B indicate the likely direction of the disordered part of EZH2^dist^ connecting the methylated peptide with the rest of the SET^dist^. (C) HMTase assays on mononucleosomes. PRC2 variants were incubated with substrate nucleosomes either un-stimulated or in the presence of 1.5 µM (+) or 15 µM (++) H3K27me3 histone H3 peptide as allosteric stimulator. (D) Stimulation of PRC2 activity by PRC2^dist^. HMTase assays on mononucleosomes were carried out with PRC2^amKR^ in the presence of PRC2^CXC^ and dimerization of automethylation-mutant variants thereof (PRC2^CXC/Δdim^ or PRC2^CXC/amKR^)

The second interaction between PRC2^dist^ and PRC2^prox^ involves the SET domain of EZH2^dist^ and EED^prox^, and has critical functional implications (Figure 3B, Figure 4A,B). The EZH2^dist^ lacks density that would correspond to the bridge helix, indicating that the am-loop containing the automethylated K510, K514 and K515 is unfolded (Figure 4B). This observation was expected, given that PRC2^dist^ is not interacting with nucleosomal DNA, and is further supported by molecular dynamics (MD) analysis (see below). The SET domain of EZH2^dist^ is positioned close to EED^prox^ such that the automethylated lysines within the unfolded am-loop of EZH2^dist^ can easily reach the allosteric methyl-lysine binding pocked of EED^prox^ (Figure 3B). Indeed, local refinement shows clear density bound at the allosteric site of the EED^prox^ (Figures 3B, 4A). We conclude that this density corresponds to the am-loop of EZH2^dist^, which is the only source of methylated peptide in our sample. Accordingly, EZH2^prox^ shows an ordered SRM helix and a bent SANT binding domain (SBD) helix, two structural hallmarks of EED-mediated activation,^3,6^ showing that PRC2^prox^ is in an activated state (Figure 4A). In contrast, neither EED^dist^ nor EED of the single PRC2-nucleosome structure reconstructed from the same dataset (which serves as an internal control), show density at the methyl-lysine binding site (Figure 4B,C). Accordingly, the SRM helix is absent from these reconstructions (and thus unfolded) and the SBD helix is in its extended conformation. Thus, while biochemically indistinguishable, PRC2^prox^ and PRC2^dist^ are in two distinct conformational and functional states: in PRC2^prox^ the bridge helix and SRM are both folded through their interactions with nucleosomal DNA and the occupied allosteric site of EED, respectively; in PRC2^dist^ those interactions are absent and thus both elements are unfolded (Figure 4A,B).

**Figure 4.**
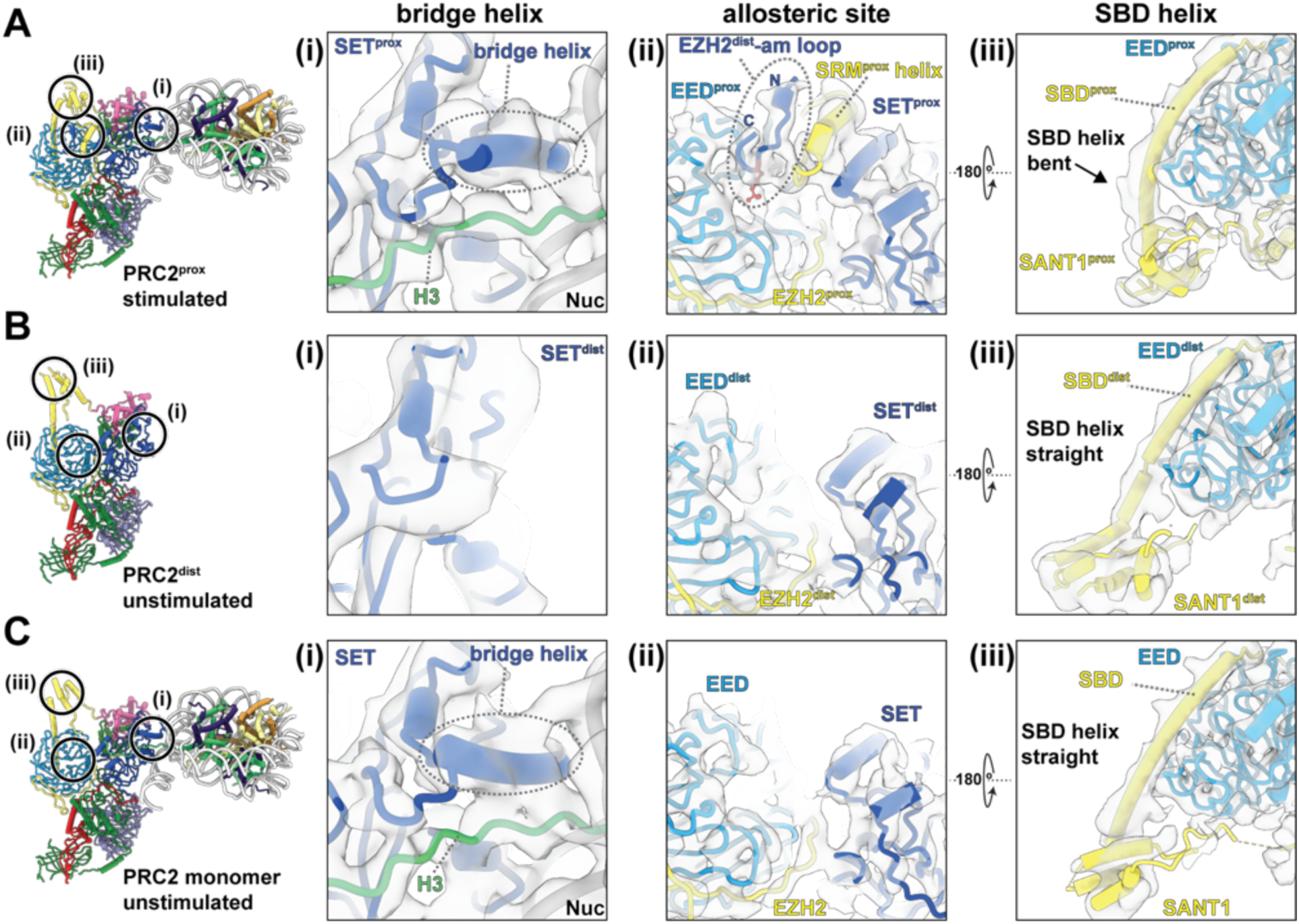
PRC2^dist^ allosterically activates PRC2^prox^ via its automethylated loop. Close-ups showing: (i) the presence or absence of the bridge helix; (ii) the occupancy of the allosteric binding site on EED and the presence or absence of the SRM helix; and (iii) the conformation of the SBD helix for PRC2^prox^(A), PRC2^dist^ (B), and PRC2 monomer (C). Only PRC2^prox^ is in an allosterically stimulated conformation, while PRC2^dist^ and PRC2 monomer are unstimulated.

### Automethylation is required for *trans*-autoactivation of PRC2

Our structural data shows that an automethylated PRC2 can serve as an allosteric activator in *trans* of a substrate-bound PRC2, inducing the hallmark structural features of allosterically activated EZH2. To investigate the functional implications of *trans* activation, we generated a series of separation of function mutants. First, in PRC2^amKR^ we mutated the am-loop lysines to arginines to mimic a non-methylated am-loop (K510R, K514R, K515R) and saw that this mutant exhibits similar activity to the wildtype PRC2 complex, as previously described.^14^ Notice that, unlike an unmethylated wild-type loop, this mutant am-loop cannot bind the EZH2 active site as a substrate. In a second mutant, termed PRC2^Δdim^ hereafter, SUZ12 lacks the short helix E_542_FLESED_548_ involved in the SUZ12-SUZ12 dimer contacts (Figure 3A). Both PRC2 variants, PRC2^amKR^ and PRC2^Δdim^, did not display any defects in overall structure, nucleosome binding, or HMTase activity upon allosteric stimulation via H3K27me3 (Figure S6A-C), indicating that automethylation and dimerization are not required for PRC2 activity when it is stimulated by an excess of an alternative allosteric activator (Figure 3C). *In vitro* HMTase activity assays to analyze allosteric activation mediated by PRC2 dimerization are complicated by the fact that PRC2 generates its own allosteric activator, i.e. H3K27me3. To address this, we made use of a third separation of function mutant that carries mutations in the CXC domain of EZH2 (K568A/Q570A/K574A/Q575A) that disrupt nucleosome engagement and abrogate HMTase activity on nucleosomes.^8,21^ However, within a dimer, this CXC mutant PRC2 (PRC2^CXC^) should still be able to adopt the distal position and serve as a *trans* activator for a PRC2^prox^.

Simultaneously, we used the PRC2^amKR^ mutant as the nucleosome modifying enzyme, which cannot itself serve as an allosteric activator. In agreement with our model, PRC2^CXC^ alone exhibited undetectable activity, but increased the amount of H3K27me3 generated by PRC2^amKR^ by ∼50% (Figure 3D and Figure S6D). Thus, although PRC2^CXC^ cannot methylate nucleosomes, it can stimulate HMTase-competent PRC2. Either disrupting automethylation or dimerization by combining the CXC and am-loop (PRC2^CXC/amKR^) or SUZ12 dimer interface mutations (PRC2^CXC/Δdim^), abrogated HMTase stimulation (Figure 3D and Figure S6D). These results agree with a model in which automethylation via dimerization results in *trans* auto-activation of PRC2.

### Impact of automethylation on bridge helix folding and nucleosome engagement

In addition to the *trans* regulatory effect exerted by means of PRC2 dimerization, automethylation could act in *cis* by affecting the stability and conformational dynamics of the bridge helix, with potential effects on substrate nucleosome binding and histone tail engagement. To analyze the relationship of automethylation and bridge helix folding, we performed molecular dynamics (MD) simulations for two distinct scenarios: (1) no methylation and (2) tri-methylation of K510, K514, K515, both in the presence and absence of nucleosome. Consistent with the disorder-to-order transition of the bridge helix indicated by cryo-EM experiments^8,16^ our simulations show that the helix is more stable when the EZH2 SET domain is nucleosome-bound (Figure 5A-C) as compared to the unbound scenario (Figure 5D-E). All simulations show an initial decrease in helicity for the bridge helix, but an overall higher degree of helicity is maintained for nucleosome-bound cases (Figure 5F). Variability between technical replicates, however, indicates dynamic conformational behavior of the bridge helix in all setups that is consistent with minimal influence of methylation on the stability of the bridge helix (Figure 5F). On the other hand, simulation data suggest that automethylation may increase the overall probability of the SET domain to interact with the H3 tail, potentially facilitating substrate engagement (Figure 5G). Overall, our MD simulations support bridge helix stabilization upon nucleosome binding, but do not show a significant effect of automethylation on bridge helix folding, and only a small effect on H3 tail engagement. Thus, automethylation likely does not have a major impact on PRC2 function in *cis* by altering the dynamics of the bridge helix.

**Figure 5:**
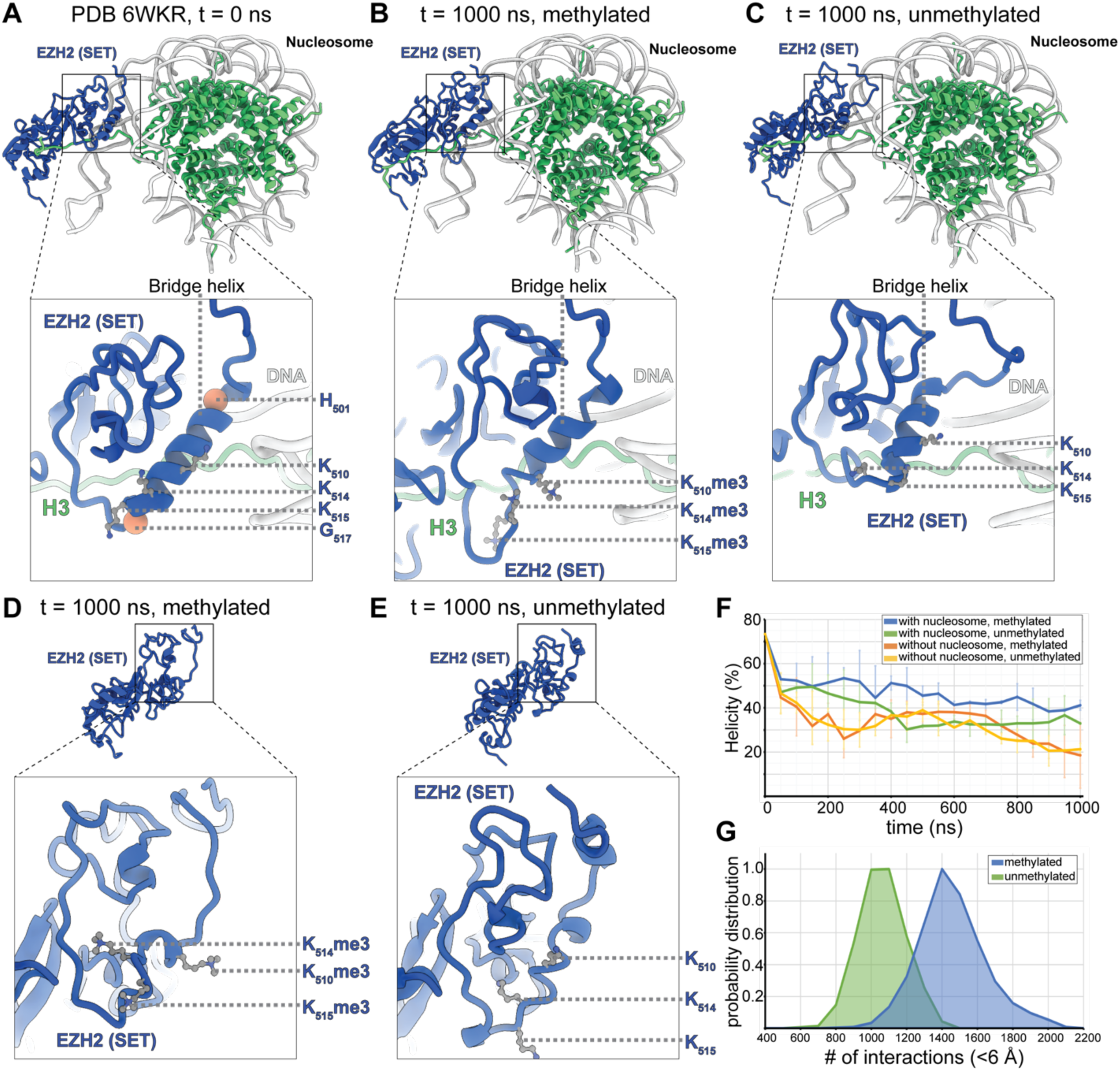
MD simulations of the EZH2 SET domain in complex with the nucleosome, analyzing the impact of auto-methylation on the conformational dynamics of the bridge helix. (A-C) Representative simulation snapshots of EZH2 SET domain bound to a substrate nucleosome at t = 0 (A), and at t = 1000 ns with the am-loop lysines K510, K514, and K515 trimethylated (B) or unmethylated (C). t = 0 corresponds to PDB 6KWR. Blue: EZH2 SET domain, green: histone proteins. The three automethylated lysines are shown in ball and stick representations. H501 and G517 are shown as coral spheres. (D-E) Snapshot of EZH2 SET domain simulation performed in the absence of nucleosome at t = 1000 ns with the am-loop trimethylated (D) or unmethylated (E). Each simulation has been performed in two replicates. (F) Analysis of the helicity of the bridge helix, including residues H501-G517 over a simulation period of 1000 ns. Helicity has been calculated as average from two technical replicates with error bars showing standard deviation. (G) Analysis of low distance (<6 Å) contacts between the H3 tail and the SET domain accumulated over the course of the 1000 ns simulation for methylated (blue) or unmethylated (green) bridge helix. Data is averaged from two experiments each.

### Role of automethylation in PRC2 gene silencing function

Based on our cryo-EM and MD results, we expect that disruption of automethylation should impact PRC2 function in transcriptional regulation by interfering with *trans*-activation. To test this hypothesis, we performed rescue experiments in EZH1/2 double knock-out murine embryonic stem cells (mESCs) in which the expression of either WT EZH2 or automethyl- mutant EZH2^amKR^ were stably restored. Restored expression of either WT or PRC2^amKR^ in our system led to comparable bulk levels of H3K27me3 (Figure 6A). In the context of retinoic acid (RA) induced differentiation along the neuronal trajectory, mESCs expressing either WT PRC2 or PRC2^amKR^ were capable of lineage commitment without evidence of a delay in differentiation (Figure 6B,D). But while the transcriptional landscape upon RA-induced differentiation was similar for the WT PRC2 and PRC2^amKR^ rescue for many genes (Figure 6C), there was a subset of genes for which the expression of PRC2^amKR^ exhibited transcriptional regulation that more closely resembled that of the EZH1/2 dKO (Figure 6C,E). This result indicates that the lack of am-loop methylation affects a subset of transcripts rather than disrupting transcriptional dynamics more globally, as would be expected for a *cis*-activating mechanism that would affect every PRC2 complex. Thus, our findings indicate a context-specific role of automethylation in which *trans*-autoactivation occurs at locations where activation cannot happen via pre-existing H3K27me3 or methylated JARID2.

**Figure 6:**
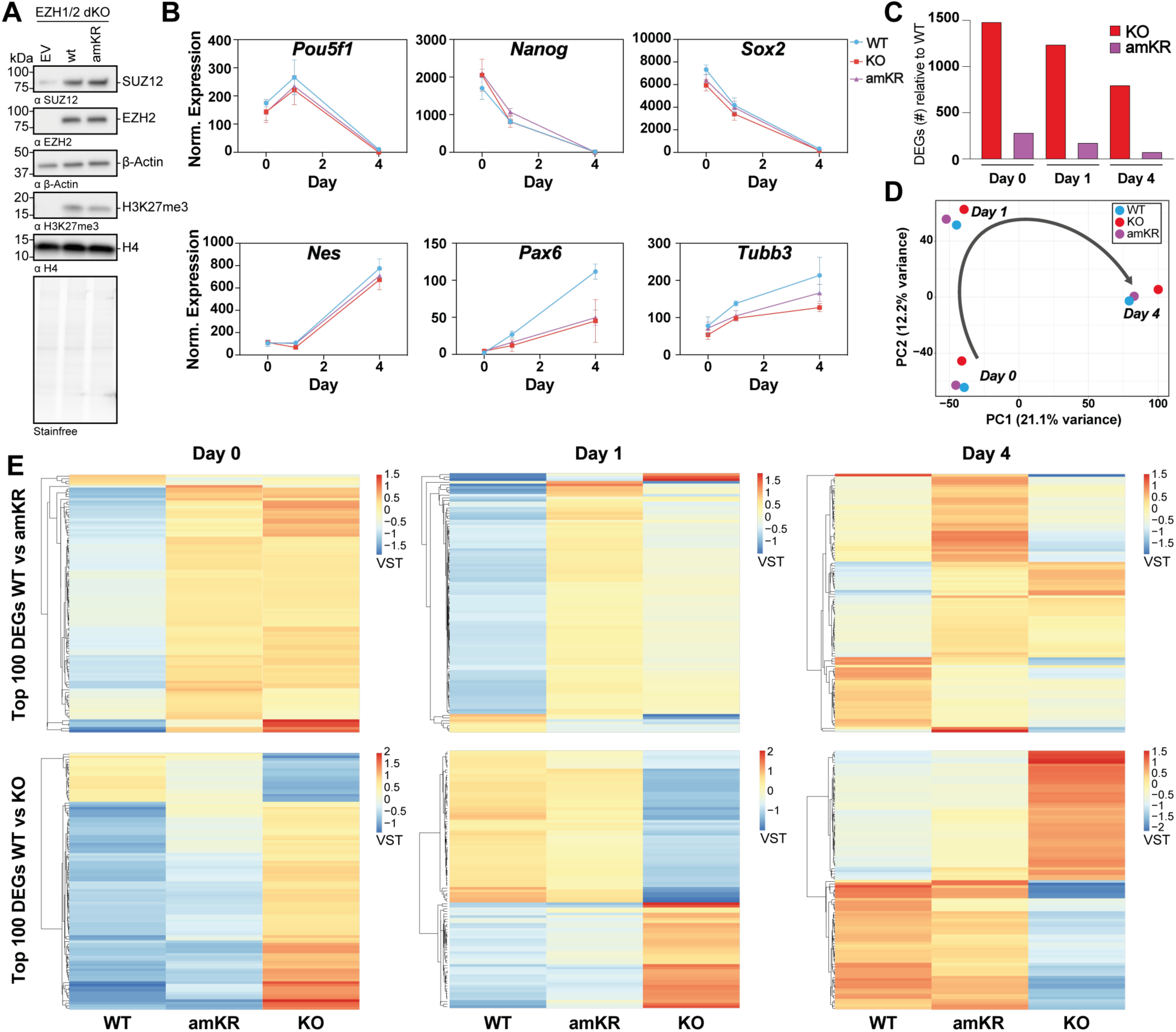
Impact of PRC2 automethylation on transcriptional dynamics during RA-induced differentiation of mESCs. (A) Western blot using the indicated antibodies of whole cell lysates obtained from EZH1/2 dKO mESCs stably transfected with either an empty vector as a negative control, the EZH2 WT, or EZH2^amKR^. (B) Gene expression profiles during RA-driven differentiation of pluripotency markers (upper panel) and neural trajectory markers (lower panel). Normalized expression data was derived from RNA-seq data and is shown as mean with standard deviation from three independent biological replicates. Samples were taken at day 0 (representing ESCs), day 1, and day 4. Blue, red and purple graphs correspond to WT, KO and amKR, respectively. (C) Number of differentially expressed genes (DEGs) relative to WT on day 0, 1, and 4, respectively. (D) PCA plot of RNA-seq data during RA-driven differentiation. Data shown as mean of three independent biological replicates. Blue, red and purple data points correspond to WT, KO and amKR, respectively. (E) Heat maps of the top 100 differentially expressed genes between WT and amKR (top panel) or WT and KO (lower panel) for day 0, day 1, and day 4, respectively. Variant stabilisation transformed (VST) data shown as a mean of three independent biological replicates.

### Chromatin context of *trans* autoactivation

We propose that different contexts will involve alternative modes of PRC2 activation via EED. The subunit composition of variant PRC2 complexes is known to affect the targeting and function of PRC2 and constitutes an important layer of its regulation. For example, JARID2 recruitment to chromatin, via the engagement of mono-ubiquitinated histone H2A^16^ or long non-coding RNAs,^22^ is a possible mechanism to specify *de novo* H3K27 trimethylation in the genome. Interestingly, we were not able to detect nucleosome-bound PRC2 dimers when JARID2 was present in the complex, even when using a JARID2 construct that lacked its lysine 116 methylation site (PRC2_J119-450_) that would otherwise compete with PRC2^dist^ binding to EED. Comparison of different PRC2 structures shows that even in the absence of K116me3, JARID2 likely outcompetes PRC2^dist^ by sterically blocking the SUZ12-SUZ12 dimerization interface, and/or impeding linker DNA binding by PRC2^dist^ (Figure S7A). Similarly, dinucleosome engagement of PRC2 representing an H3K27me3 spreading site, in which an H3K27me3- bearing nucleosome allosterically activates EZH2 to methylate an adjacent nucleosome via EED,^8^ is sterically incompatible with dimerization as shown here. In a hetero-dinucleosome, the methylated nucleosome would clash with the SANT2 domain of the PRC2^dist^ (Figure S7B). Thus, allosteric *trans*-activation by dimerization is a context-specific mechanism of PRC2 activation that occurs as an alternative to activation by JARID2 K116me3 or H3K27me3 bearing nucleosomes.

The linker histone H1 is another determinant of chromatin context that has been proposed to cooperate with PRC2 to suppress gene expression through an unknown mechanism.^23^ Interestingly, cryo-EM analyses of PRC2 incubated with H1 bearing nucleosomes suggest that H1 binding and PRC2 dimerization are mutually exclusive, since all reconstructions of PRC2 dimers lacked density for H1 (Figure S8A). On the other hand, we could see clear EM density corresponding to H1 when a PRC2 variant containing JARID2_119-450_ was used, a condition that prevents PRC2 dimerization (Figure S8B,C). Notice that this PRC2_J119-450_ is seen in an inactive conformation, as expected given the absence of an allosterically activating methylated peptide, and also that there is no direct interaction between H1 and PRC2_J119-450_. Comparison of the nucleosome-H1, nucleosome-H1- PRC2_J119-450_ and nucleosome-PRC2^prox^-PRC2^dist^ structures shows that H1 binding gives rise to a linker DNA trajectory that is not compatible with PRC2^dist^ binding (Figure S8D,E). Therefore, these observations indicate that genomic occupancy of H1 is incompatible with the *trans*-autoactivation of PRC2 by dimerization.

## Discussion

### An extended model for *de novo* establishment of H3K27me3

The activity of the PRC2 chromatin regulator is critical in development to both establish and maintain cell identity. Key to its function are the regulation of its genomic targeting and the local regulation of its HMTase activity. The discovery of the stimulatory effect of EZH2 automethylation on HMTase activity^14,15^ led us to hypothesize that automethylated EZH2 may act via the well-established allosteric methyl-lysine binding site in the regulatory subunit EED. To investigate the underlying mechanism, we studied the impact of this modification on PRC2 conformation and nucleosome engagement in the absence of any other methylated peptide. Our studies led us to discover a chromatin-dependent PRC2 dimer, in which the automethylated am- loop of a PRC2^dist^ binds the allosteric site in EED^prox^ of a PRC2^prox^ that is engaged with the tail of the substrate nucleosome (Figure 7). The two PRC2 complexes are present in two distinct conformational states, with only PRC2^prox^ showing the structural hallmarks of allosteric activation (a bent SBD helix and stabilized SRM). In agreement with the model of PRC2 regulation that emerges from these structural observations, functional assays using separation of function mutants show that nucleosome-binding deficient PRC2 (PRC2^CXC^) can serve as an allosteric activator, as long as automethylation and dimerization are not impaired.

**Figure 7.**
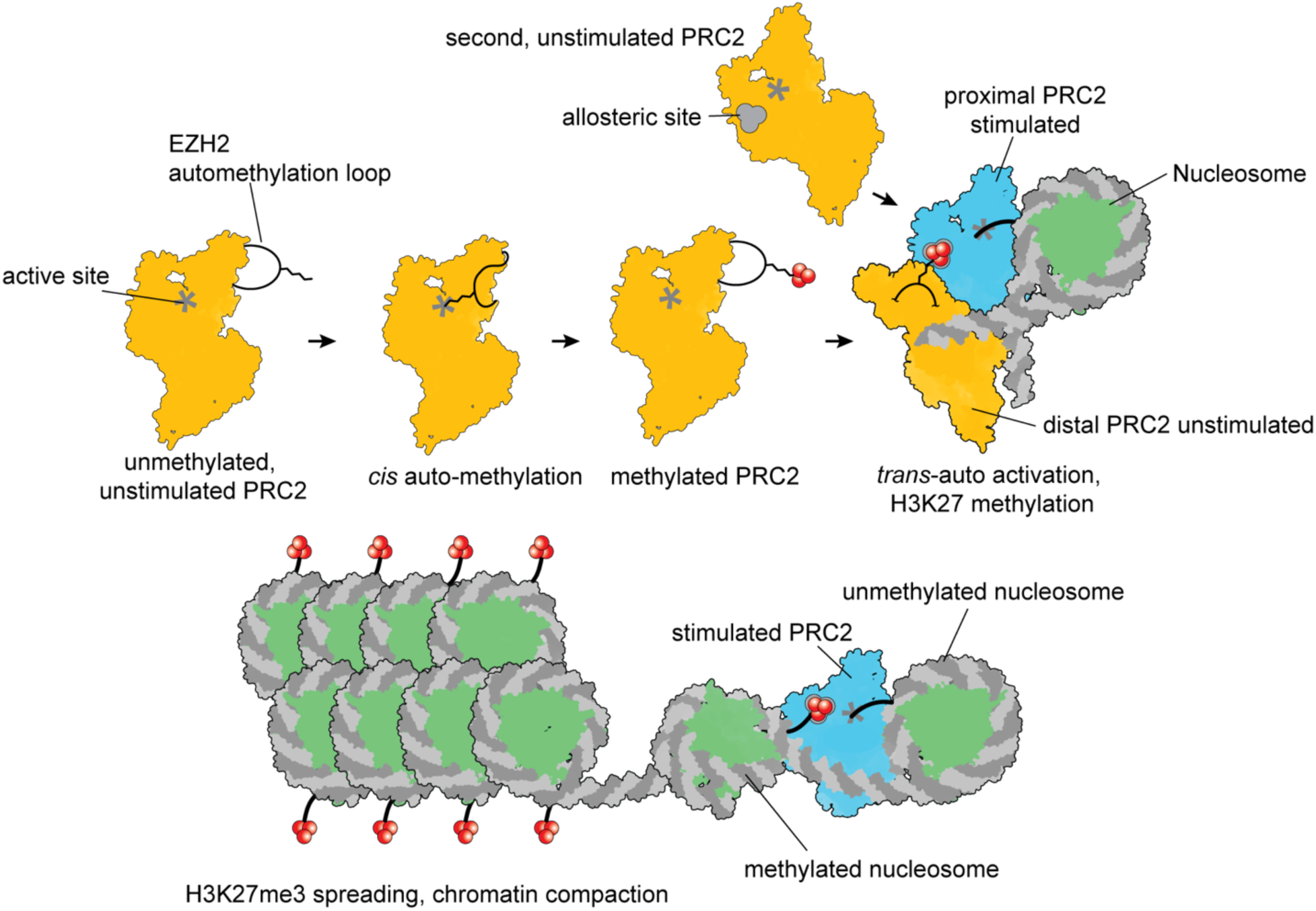
Trans-auto activation model. PRC2 that undergoes automethylation in cis can act as an allosteric activator in trans for a second PRC2 that then methylates H3 on the nucleosome as they interact with each other in the context of chromatin. Once nucleosomes containing H3K27me3 accumulate, PRC2 can be allosterically activated to further spread the H3K27me3 mark and ultimately cause chromatin compaction. We propose that trans-auto activation of PRC2 enables initiation of H3K27me3 domains in the absence of other stimulating cofactors.

The *trans* activation mechanism shown here employs the established EED-EZH2 allosteric communication axis that has been well studied for other activators, i.e. H3K27me3 and JARID2 K116me3. In all three cases, the methylated peptide binds EED and causes the stabilization of the SRM and the active conformation of the EZH2 SET domain, underscoring the central role of this mechanism in PRC2 regulation. Therefore, it is expected that the activation by all three established EED ligands represent alternative pathways of PRC2 activation. In agreement with this notion, the *in vitro* HMTase activity of automethylation or dimerization mutant PRC2 is unaffected when stimulated by an excess of H3K27me3 peptide. Additionally, we show that *trans*-autoactivation by dimerization and activation by JARID2 K116me3 or H3K27me3 are mutually exclusive, since H3K27me3 bearing nucleosomes would clash with PRC2^dist^, and no PRC2 dimers were observed when JARID2 containing PRC2 variants were used. We conclude that dimerization-mediated auto-activation is an EED-dependent mechanism of PRC2 activation that occurs at genomic loci in which automethylated EZH2 is the only available activator of PRC2. This model is supported by our transcriptomics analyses in mESCs showing that the abrogation of automethylation affects the regulation of a subset of transcripts upon cellular differentiation, while global H3K27me3 levels remain largely unaffected. Accordingly, previously published rescue experiments using am-loop mutant PRC2 showed reduced H3K27me3 levels in transient rescue experiments,^14,15^ but not when am-loop mutant EZH2 was expressed for longer time periods.^15^ We conclude that automethylation-mutant EZH2 fails to establish PRC2 function in distinct genomic contexts rather than globally. This “specificity” is incompatible with a strong *cis* regulatory effect of EZH2 automethylation, which would affect PRC2 activity independent of context. In agreement with the absence of a strong *cis* regulatory mechanism of automethylation that could affect bridge helix folding and/or substrate nucleosome engagement, our MD analyses show minimal effects of automethylation on bridge helix folding and only a limited impact on histone tail engagement by PRC2. Accordingly, nucleosome binding was not strongly affected by am-loop mutations in PRC2^amKR^ and previous work showed that automethylation-mutant PRC2 does not show a defect in chromatin engagement and genome wide occupancy.^14^

It has been proposed that the unmethylated am-loop could bind the active site of EZH2 competing with other substrates, and thus, am-loop methylation could release an auto-inhibited state of PRC2.^15^ In the context of this *in cis* hypothesis, the PRC2^amKR^ mutant simulates a state in which the auto-inhibition is released, thus resembling the methylated am-loop. Under this assumption, one would expect increased activity of PRC2^amKR^, which is not observed. Such a hypothesis also cannot explain the stimulation by PRC2^CXC^ of PRC2^amKR^. While we cannot exclude a possible impact of am-loop binding to the EZH2 active site on PRC2 activity, our work indicates that mechanisms acting in *cis* alone are insufficient to explain the stimulatory effect of EZH2 automethylation on PRC2 activity.

In a physiological context, the *de novo* establishment of H3K27me3 can be triggered via PRC2 activation by methylation of JARID2 K116, which is recruited to genomic loci via H2A K119ub^16,24^ or long non-coding RNAs.^5,22^ We propose that PRC2 dimerization could initiate H3K27me3 in the absence of JARID2, e.g. in cells that lack JARID2 expression or at genomic loci that do not recruit JARID2. Unlike activation by JARID2, dimerization requires two PRC2 complexes, thus higher local PRC2 concentration. This likely implicates local concentration levels of PRC2 in regulating initial H3K27me3 deposition, e.g. involving factors that recruit but not themselves activate PRC2. Further context specificity is suggested by our observation that histone H1 binding and *trans* autoactivation by dimerization are incompatible. These findings underscore the intricate ways in which chromatin regulators integrate cues from the local chromatin environment for their targeted and regulated function. Our work showcases the central role that the regulatory EED subunit has in instructing PRC2 HMTase activity in diverse contexts. Moreover, *trans* autoactivation by dimerization of PRC2 shows that mechanisms that integrate auto-catalysis, homo-oligomerization and allosteric regulation are not limited to the classical examples of kinase autophosphorylation,^9,10^ but are highly relevant to multi-protein complexes that regulate chromatin function via histone modification.

Homotypic interactions of enzymes that are regulated through auto-catalysis can have various functional implications, including the amplification of regulatory signals, impact on substrate recognition, and additional layers of regulation, such as feedback loops or temporal control. EZH2 automethylation likely enables the *de novo* deposition of H3K27me3 in distinct chromatin contexts. One could even imagine that the stable methylation of lysines in the am-loop of EZH2 may act as a molecular memory of PRC2 activity, potentially enabling *trans* activation of multiple complexes. Furthermore, the system may be further fine-tuned by combining different variants of PRC2 within one dimer. For example, EZH1-containing complexes, which show little HMTase activity themselves, could potentially serve as allosteric activators in hetero- dimers of PRC2/EZH1 and PRC2/EZH2, since the am-loop is conserved between the two proteins. The overall ability of additional cofactors and subunit variants to affect PRC2 targeting and to facilitate or impede the formation of PRC2 dimers remains to be investigated. Another open question is whether and how automethylation, a prerequisite for dimerization-mediated activation, is itself regulated. Thus, automethylation and allosteric dimerization add further layers of complexity to PRC2 targeting and regulation, and provide support for the critical role played by trimethyl-lysine binding to the regulatory subunit EED. Future work will determine whether regulation by dimerization extends to other key epigenetic factors that have been shown to auto-modify.^11–13^

## Acknowledgements

We thank Dr. Anthony Iavarone of the QB3/Chemistry Mass Spectrometry Facility at UC Berkeley for assistance with mass spectrometry measurements, A. Chintangal, K. Stine and P. Tobias for computational support, Dr. D. Toso at the Cal Cryo facility for support with cryo-EM data collection, Dr. R. Glaeser and Dr. B.-G. Han for advice concerning streptavidin grid preparation, P. Rodewald for support with protein and negative-stain grid preparation. We thank Dr. E. Behrmann and Dr. Monika Gunkel of the StruBiTEM cryo-EM facility at the University of Cologne for support, as well as Dr. A. Schauss and Dr. F. Gaedke of the imaging facility at the CECAD, Cologne, these facilities were used for negative-stain and cryo-EM sample screening and data collection. We thank Dr. D. Pasini for kindly sharing WT, SUZ12 KO and EZH1/2 dKO cell lines. We thank Dr. A. Skoultchi and Dr. Sean Healton for providing the H1 protein and protocols. We thank Dr. V. Kasinath for discussion of the data and feedback. We acknowledge the Cologne Center for Genomics for its support of RNA-seq library preparation and sequencing. We thank the Regional Computing Centre (RRZK) of the University of Cologne for its support.

Molecular graphics and analyses performed with UCSF ChimeraX, developed by the Resource for Biocomputing, Visualization, and Informatics at the University of California, San Francisco, with support from National Institutes of Health R01-GM129325 and the Office of Cyber Infrastructure and Computational Biology, National Institute of Allergy and Infectious Diseases.

## Funding

EP, JR, LCZ and SP are funded by CMMC core funding (JRG XI). SP and EP are supported by the Deutsche Forschungsgemeinschaft (DFG, German Research Foundation) - SFB1430 - Project-ID 424228829, and the CANTAR network funded by the Ministry of Culture and Science of the state of Northrine-Westphalia.

EN is funded by the National Institutes of Health, NIGMS grant GM127018. She is a Howard Hughes Medical Institute Investigator.

PH, MNH, OvR and RHH are supported by the DFG CRC1399, DFG RU5504, DFG (HA 8562/4-1), CANTAR, CMMC core funding (JRG X) and the Fritz-Thyssen Foundation.

TC is supported by the National Science Foundation Graduate Research Fellowship under grant number DGE 2146752.

AS and KYS were supported by Los Alamos National Laboratory (LANL) Laboratory Directed Research and Development grant 20210082DR and LANL Laboratory Directed Research and Development grant 20210134ER. AS and KYS acknowledge generous allocations of computational resources on the Chicoma supercomputer by Los Alamos National Laboratory Institutional Computing.

## Author contributions

PVS and EP designed and performed experiments, interpreted results. EN and SP designed and supervised the project and interpreted results. TC helped with the sample preparation, performed experiments, interpreted results. LCZ and JR performed experiments. RHH designed and supervised transcriptomics experiments, PH performed transcriptomics data analysis, interpreted results, and created figures, MNH performed data analysis, OVR prepared samples for transcriptomics. AS performed MD simulation an analysis. AS and KYS interpreted MD results. PVS, EP, SP, TC and EN wrote the manuscript with input from all authors.

## Competing interests

The authors declare no competing interests.

## Data and materials availability

Cryo-EM density maps and fitted models have been deposited in the Electron Microscopy Data Bank (EMDB) and the Protein Data Bank (PDB) under the accession numbers EMD-41110 and PDB 8T9G for the PRC2 dimer bound to nucleosome, EMD-41141 and PDB 8TAS for the PRC2 monomer bound to nucleosome, EMD-41146 and PDB 8TB9 for PRC2_J119-450_ bound to H1-nucleosome and EMD-41147 for the H1-nucleosome complex. All cryo-EM micrographs used for this study were deposited in the Electron microscopy public archive (EMPIAR) under the accession code EMPIAR- 11607. The code used in this study to subtract the streptavidin lattice from the electron micrographs is available on github under https://github.com/pvsauer/StreptavidinLatticeSubtraction. Raw gel images are shown in Figure S9.The transcriptomics data from this study is available under GEO record identifier GSE234793.

## Methods

### Cloning, expression, and purification of PRC2

PRC2 was cloned, expressed, and purified as previously described.^8,16^ Briefly, full length sequences of EZH2 isoform 2, EED, RBAP48, strep-tagged AEBP2 and residues 80-685 of SUZ12 (residues 80-685) were cloned into the MacroBac system for baculovirus expression in HighFive insect cells.^25^ For experiments involving the subunit JARID2, residues 119-450 (excluding the methylated K116 residue) of JARID2 were also included in the MacroBac plasmid. Expression of PRC2 occurred for 72 hours at 27 °C and pellets were stored at -80 °C until use. All purification steps were performed at 4°C. Pellets were resuspended in 25 mM HEPES, pH 7.9, 250 mM NaCl, 5% glycerol, 0.1% NP-40, 1 mM TCEP, supplemented with 10 μM leupeptin, 0.2 mM PMSF, protease inhibitor cocktail (Roche) and benzonase (Sigma-Aldrich). Cells were lysed by sonication and cleared by centrifugation at 35 000g for 45 minutes. The supernatant was incubated with Step-Tactin Superflow Plus resin (Qiagen) for 6 hours and then washed with low (25 mM HEPES, pH 7.9, 150 mM NaCl, 1 mM TCEP, 5% glycerol, 0.01% NP40) and high salt buffers (25 mM HEPES, pH 7.9, 1 M NaCl, 1 mM TCEP, 5% glycerol, 0.01% NP40) followed by elution with 10 mM desthiobiotin.

The eluate was pooled and incubated with TEV protease over night to cleave off the affinity tag, followed by size exclusion chromatography using a Superose 6 3.2/300 column (Cytiva) equilibrated with 25 mM HEPES pH 7.9, 150 mM NaCl, 2 mM MgCl2, 10% glycerol, and 1 mM TCEP. Purified complex was flash frozen in liquid nitrogen and stored at −80 °C as single-use aliquots.

For the HMTase activity assays, the EZH2 and SUZ12 mutations required for the PRC2 mutants were introduced by site directed mutagenesis, and the subunits were assembled into multi-gene plasmids for baculoviral expression using the GoldenBac assembly protocol.^26^ These mutants were expressed in *T. ni* insect cells (Expression Systems) and purified as described above.

### Nucleosome purification

Xenopus histones (H2A, H2B, H3, and H4) were expressed and purified as described previously.^27^ The nucleosomal DNA contains a CpG Island sequence and a 5′ biotin tag and was assembled by large scale PCR, purified over an anion exchange column, and further purified by ethanol precipitation. The 226-base pair (bp) nucleosome DNA sequence used for all the studies with Xenopus nucleosomes was 5′ (biotin) CACGCGACTGTGTGCCCGTCAGACGCTGCGCTGCCGGCGG<u>ctggagaatcccggtgccgaggccgc</u> <u>tcaattggtcgtagacagctctagcaccgcttaaacgcacgtacgcgctgtcccccgcgttttaaccgccaaggggattactccctagtctccaggcacgtgtcagatatatacatcctg</u>tatgcatgcatatcattcgatcggagctcccgatcgatgc - 3′. The CG-rich sequence used is capitalized and the 601-nucleosome positioning sequence^28^ is underlined. For nucleosome assembly, equimolar amounts of all histones were dialyzed into histone refolding buffer (2 M NaCl, 10 mM TRIS, 5 mM EDTA), and the octamer was purified using a Superdex 200 10/300 size exclusion column (Cytiva). The DNA and octamer were mixed in a 1:1.1 ratio and purified over a BioRad prep cell after overnight gradient salt dialysis, as described previously.^27^

To create H1-containing nucleosomes, His-tagged xenopus histone H1.0 and His-tagged murine nuclear assembly factor 1 (mNAP1) were cloned into and recombinantly expressed in BL21-DE3 *E. coli* and purified as follows: For H1, bacterial cell pellets were resuspended in lysis buffer (1 M NaCl, 20 mM Tris pH 7.4, 10 mM imidazole, 10% glycerol, 0.5 mM TCEP) supplemented with DNAse, PMSF and EDTA free protease inhibitor (Roche), before lysis by sonication. After an addition of 1% (v/v) Triton X-100 and centrifugation at 35 000 g for 30 minutes at 4°C, the clarified lysate was incubated on charged Nickel beads (Qiagen) that have been pre-equilibrated in lysis buffer containing 1% (v/v) Triton X-100. Beads were washed with 5 column volumes (CV) of lysis buffer, followed by 5 CV of wash buffer (500 mM NaCl, 20 mM Tris pH 7.4, 10 mM imidazole, 0.5 mM TCEP) until no more protein eluted as monitored by Bradford reagent. H1 was eluted with elution buffer (500 mM NaCl, 20 mM Tris pH 7.4, 500 mM imidazole, 0.5 mM TCEP) and the peak fraction collected for dialysis into 200 mM NaCl, 20 mM Hepes pH 7.4, 0.5 mM TCEP at 4°C over-night. Dialyzed protein was subjected to cation exchange chromatography using a Mono S 5/50 GL column (Cytiva) and subjected to a salt gradient to 1 M NaCl. H1 containing fractions were pooled, frozen in liquid nitrogen and stored at -80° C until further use.

For NAP1, purification was essentially carried out the same as for H1 except for following steps: no Triton X-100 was added during purification and ion exchange chromatography was carried out as anion exchange chromatography and therefore a Mono Q 5/50 GL column was used (Cytiva). Consequently, the ion exchange buffer and the preceding dialysis buffer contained 20 mM Tris pH 7.4 instead of 20 mM HEPES.

NAP1 mediated H1 deposition on nucleosomes was carried out as described previously.^29^ H1 and NAP1 were mixed in a 1:2 molar ratio and incubated at 30 C for 30 minutes in 100 mM NaCl, 20 mM Tris pH 7.4, 0.5 mM EDTA, 10 % glycerol, 1 mM DTT. NAP1-H1 complexes were then incubated with biotinylated nucleosome in a 5 Molar excess for 30 minutes at room temperature. The sample was then directly used for cryo-EM experiments as described below.

Excess NAP1-H1 was washed away from biotinylated nucleosomes during cryo-EM sample preparation using streptavidin affinity grids.

### Mass spectrometry

To analyze determine the number of methylated lysine residues on the automethylation loop, ∼150 μg of PRC2 were first unfolded and reduced in fresh 6.4 M urea and 10 mM DTT and incubated at 55 °C for 20 minutes. Cysteines were alkylated using 20 mM iodoacetamide, followed by incubation at RT for 30 min in the dark and subsequent quenching with an additional 30 mM DTT. Quenching was allowed to occur for 20 min at RT before dialysis at 4 °C against 50 mM Tris pH 7.7, 5 mM CaCl2, 2 mM EDTA and 5 mM DTT to remove urea and iodoacetamide. The sample was then digested with 500 ng Arg-C endopeptidase overnight at RT. The reaction was stopped by incubation for 10 minutes at 95 °C. Liquid chromatography – mass spectrometry measurements of the sample were performed in the QB3/Chemistry Mass Spectrometry Facility at UC Berkeley as described elsewhere.^30^ Briefly, digested PRC2 was analyzed using a Synapt G2-Si ion mobility mass spectrometer equipped with a nanoelectrospray ionization source (Waters) in line with an ACQUITY M-class ultraperformance LC system.

Raw data acquisition was controlled using MassLynx software (version 4.1), and peptide identification and relative quantification were performed using Progenesis QI for Proteomics software (version 4.0; Waters). Calculation of the percentage of lysine methylation (mono-, di-, tri-, or unmethylated) was performed by dividing the abundance of a peptide bearing one or several modifications by the total abundance and multiplying by 100.

### Negative stain EM

Negative stain analysis of PRC2 was carried out essentially as described before.^8^ Briefly, 4 µl of 200 nM PRC2 were incubated on a continuous carbon grid (EMS) for 45 sec, followed by five successive short incubation steps with 2% (wt/vol) uranyl formate. Excess stain was removed by blotting with filter paper and the grids were dried. Screening and data collection was done using a Talos L120C (Thermo Fisher Scientific) and EPU for automated data acquisition, at an electron dose of 25 e/Å^2^ and a nominal pixel size of 2.44 Å/px. Data processing was done in cryosparc,^31^ CTF estimation was done using CTFFIND4,^32^ and particle picking using the blob picker in cryosparc. For WT PRC2, PRC2^amKR^ and PRC2^Δdim^, 117,434, 248,000 and 176,329 particle images were extracted based on initial picks from and 242, 398 and 286 manually curated micrographs, respectively. Several rounds of 2D classification led to subsets of classes with typical structural features of PRC2, which were overall comparable between WT PRC2 and PRC2^amKR^ and PRC2^Δdim^ mutants. Representative, typical views of intact PRC2 were chosen and adjusted for PRC2 orientation.

### Cryo-EM grid preparation

To prevent damage of PRC2 by interactions with the air water interface we used streptavidin affinity grids manufactured in-house, as described previously.^16,33,34^

All PRC2-nucleosome complexes were assembled by incubating 200 nM biotinylated nucleosome (containing or lacking H1) with 800 nM PRC2 and 100 μM SAH in 25 mM HEPES pH 7.9, 50 mM KCl, 1 mM TCEP for 30 minutes at RT. 4 μl of the complex were incubated on rehydrated Quantifoil Au 2/2 grids containing the streptavidin affinity layer and incubated for 5 minutes in a humidified chamber. The grid was then washed with two times 10 μl of buffer containing 25 mM HEPES pH 7.9, 50 mM KCl, 1 mM TCEP, 4% Trehalose, and 0.01% NP40. Excess buffer was wicked away with filter paper before adding an additional 2.5 μl of the same buffer. After transfer of the grid into a TF Mark IV Vitrobot the grid was manually blotted for 2- 3 s at 18 °C and 100% before plunging it into liquid ethane.

### Data collection and processing

For the PRC2 dimer, two datasets (dataset 1 and 2) were collected on a FEI Titan Krios G2 cryo-electron microscope operating at 300 kV, equipped with a GIF quantum energy filter and a GATAN K2 direct electron detector in super resolution mode. 3,894 micrographs were collected for dataset 1 and 4,062 micrographs were collected for dataset 2. For each exposure a total of 40 frames were collected with a total dose of 50 e^−^/Å^2^ at a super resolution pixel size of 0.575 Å/pix while varying the defocus between -1.5 and -3.5 μm. Movies were motion corrected and dose weighted using MotionCor2 ^35^ before subtraction of the streptavidin lattice using in-house MATLAB scripts. ∼600k particles were picked using the convolutional neural network picker implemented in EMAN2.^36^ CTF estimation, particle extraction and initial rounds of 2D and 3D classification were carried out in Relion 3.0 ^37^ for initial clean-up of the particle stack. After merging particles from both datasets and another round of 3D classification a class corresponding to the PRC2 dimer bound to the nucleosome and a class corresponding to the PRC2 monomer bound to the nucleosome became apparent. Particles were transferred to Cryosparc v4.0 ^31^ for all further processing. The dimer and the monomer classes were refined independently to resolutions of 6.2 and 4.1 Å, respectively, determined according to the gold-standard FSC = 0.143 criterion.^38,39^ To overcome continuous flexibility inherent to the complexes we used 3DFlex as implemented in Cryosparc to improve the quality of our maps.^19^ Dividing the PRC2 dimer bound to nucleosome into three bodies, where both PRC2 protomers are attached to the nucleosome, and using 5 latent dimensions during the program training phase, revealed several modes of relative motion in the complex and improved the quality of the distal PRC2 protomer (Figure S4). Focused refinements with search parameters adjusted for large movements yielded the final maps which were then filtered by local resolution using manually adjusted B-factors to prevent over-sharpening. The final maps are represented in Figure S2.

For PRC2_J119-450_-H1-Nucleosome, 14,470 movies (dataset 3) were collected on a FEI Titan Krios G2 cryo-electron microscope operating at 300 kV, equipped with a GIF quantum energy filter and a GATAN K3 direct electron detector in super resolution mode. For each exposure a total of 50 frames were collected with a total dose of 50 e^−^/Å^2^ at a super resolution pixel size of 0.575 Å/pix while varying the defocus between -0.8 and -2.5 μm. After motion correction and streptavidin lattice subtraction, 3,658,724 particles were picked using Cryolo. CTF estimation, particle extraction and initial rounds of 2D and 3D classification were carried out in Relion 3.1 for initial clean-up of the particle stack. After another round of 3D classification, a class corresponding to the PRC2_J119-450_ bound to the H1-nucleosome became apparent. Particles were transferred to Cryosparc v. 4.0 and the map refined to a resolution of 4 Å, determined according to the gold-standard FSC = 0.143 criterion. Local refinements of the H1-nucleosome and PRC2_J119-450_ yielded the final maps with resolutions of 3.6 Å each which were then filtered by local resolution. For the H1.0-nucleosome complex, 2,236 micrographs were collected on a Talos Arctica electron microscope operating at 200kV and equipped with a Gatan K3 direct electron detector using a final pixelsize of 1.14 Å/pix. After motion correction, streptavidin lattice subtraction and CTF estimation data was processed in Cryosparc v4.0 using a standard workflow. A final particle set of 44,742 yielded a reconstruction with a resolution of 3.14 Å. Data was further processed by using 3DFlex to improve regions of the map suffering from flexibility.

### Model building and visualization

To obtain a model for the allosteric PRC2 dimer and the PRC2 monomer we used a trimmed model of PRC2 bound to an ubiquitylated nucleosome (pdb 6WKR^16^) as a starting point to perform flexible fitting using Isolde v1.5 in UCSF ChimeraX v1.5 ^40,41^ into locally refined maps and model building in Coot,^42^ applying appropriate model restraints. Nucleosomal linker DNA was modeled using ChimeraX and then also flexibly fitted into the density using Isolde v1.5. The automethylation loop was modeled using a fragment of JARID2 present in the input structure and also fitted using Isolde v1.5 and Coot.

The same strategy was used to obtain a model for the monomeric PRC2 bound to nucleosome and for PRC2_J119-450_ bound to H1-nucleosome. For the H1 containing nucleosome, PDB 3NL0 was used as a starting model. All models were then subjected to real space refinement in phenix v1.2,^43^ enabling local grid search, global minimization and ADP refinement, with Ramachandran restraints enabled but secondary structure restraints disabled. For modeling of the automethylation loop, the visually most likely sequence was modeled into the density (with EZH2 K514 being recognized by EED) and refined as described. To remove author bias, all am-loop residues except trimethylated lysine were then mutated to alanine in Coot, renamed to UNK and subjected to another round of ADP-only refinement in phenix. All final models were created by combining the local models into a composite model and refining the final models against the respective consensus reconstruction with model restraints enabled. Refinement parameters and model validation parameters are reported in Suppl. Table 1. ChimeraX v1.5 ^40^ was used to visualize maps and models.

### Histone methyltransferase (HMTase) assay

To perform the HMTase assay, reactions were carried out in a total volume of 12 µL containing 200 nM nucleosome and 400 nM PRC2 in a reaction buffer (25 mM HEPES pH 7.9, 50 mM NaCl, 2.5 mM MgCl2, 0.25 mM EDTA, 5% glycerol, 1 mM DTT and 80 µM SAM).

Prior to adding the nucleosome, the reaction mix was preincubated at room temperature for 1 hour to allow for automethylation. The reaction proceeded at room temperature for 90 minutes and was quenched by the addition of 5x loading buffer and heat inactivation at 95°C for 5 minutes. In case of peptide stimulation, the peptides were added immediately after the nucleosome. Separation by gel electrophoresis was performed with 4–20% Mini-PROTEAN® TGX™ precast protein gels (BioRad) and the stain-free signal was detected according to the manufacturer’s instructions. Proteins were subsequently transferred to a 0.2 µM PVDF membrane at 90V for 10 min and 60 V for 30 min. The membranes were probed with antibodies against H3K27me3 (Active Motif, 39155) and H4 (Cell Signaling, L64C1). Reactions were performed in multiplets and detected with a ChemiDoc MP (BioRad). Densitometric analysis was performed using Image Lab Software version 6.1.0 (BioRad) by background-correcting the signal to the negative control and normalizing it against the WT signal. GraphPad Prism was used for visualization.

### Electrophoretic mobility shift assay (EMSA)

EMSA was performed using a 5% native TBE gel in 0.2x TBE buffer with a total volume of 15 µL reactions of 50 µM nucleosome and increasing concentrations of PRC2 in triplicates in binding buffer (25 mM HEPES pH 7.9, 50 mM NaCl, 1 mM DTT and 100 µM SAH). The reaction mixture was incubated at room temperature for 30 minutes to allow for binding. The gels were then stained with SYBR™ Gold (Thermo Fisher) according to the manufacturer’s instructions. The stained gels were imaged using a ChemiDoc MP (BioRad) imager, and densitometric analysis was performed using Image Lab Software version 6.1.0 (BioRad). The bands of the shifted (bound) and free nucleosomes (unbound) were identified and boxed out.

After background correction, the bound signal was divided by the sum of both signals to determine the bound fraction.

### Murine embryonic stem cell cultivation and differentiation

The EZH1/2 dKO mESCs (N/A Strain of origin 129P2/Ola) used in this study were obtained from the Pasini lab and previously characterized.^44^ The cells were cultured on 0.1% gelatin-coated dishes in mESC media consisting of GMEM (Gibco) supplemented with 20% ES-grade fetal bovine serum (Gibco), 2 mM glutamine (Gibco), 100 U/ml penicillin, 0.1 mg/ml streptomycin (Gibco), 0.1 mM non-essential amino acids (Gibco), 1 mM sodium pyruvate (Gibco), 50 µM ß-mercaptoethanol (Gibco), 1000 U/ml ESGRO Leukemia Inhibitory Factor (LIF, Sigma Aldrich, ESG1107), and GSK3β and MEK 1/2 inhibitors (Axon Medchem BV) to a final concentration of 3 μM and 1 μM, respectively. For maintaining a confluency of between 60 and 70%, cells were passaged every 2–3 days by washing twice with phosphate-buffered saline (PBS) and dissociation with 0.25% Trypsin (Life Technologies, 25200056).

For transfection, EZH2 wild type and EZH2amKR were cloned into a pPB_PGK plasmid and co-transfected with a piggyback transposase using Lipofectamine 2000 (Thermo Fisher Scientific) following the manufacturer’s instructions and were selected with puromycin (1 µg/ml).

For differentiation mESCs were seeded at a density of 10500 cells/cm^2^ in mESC media lacking LIF, GSK3β, and MEK 1/2 for 12 hours to allow for cell attachment. The media was then exchanged and supplemented with 0.1 µM all-trans-retinoic acid, and subsequently changed every 48 hours.

### Whole cell lysis and western blotting

Total protein lysis was performed by incubating the cells on ice for 30 minutes followed by sonication in ice-cold RIPA buffer (50 mM Tris-HCl pH 7.4, 150 mM NaCl, 1 mM EDTA, 1 % NP-40, 1% Na-deoxycholate, 0.1 % SDS) supplemented with protease inhibitors and 1 µg/mL Benzonase (produced in-house). Protein concentration was determined with Pierce™ Rapid Gold BCA Protein-Assay-Kit (Thermo Fisher) and normalized to 60 µg before being supplemented with Laemmli sample buffer. Protein lysates were separated via SDS-PAGE and transferred to PVDF membrane at 90V for 120 minutes. The membranes were probed with antibodies against H3K27me3 (ActiveMotif, 39155), SUZ12 (Cell Signaling, 3737S), EZH2 (Cell Signaling, 3147S), H4 (Cell Signaling, L64C1), and β-Actin (Sigma-Aldrich, A5441-.2M). The proteins were detected with a ChemiDoc MP (BioRad)

### RNA isolation and RNA seq

Total RNA was isolated from cells using NucleoSpin RNA Kit (MACHEREY NAGEL, cat. no. 740955) according to manufacturer’s protocol. Libraries for RNA seq were generated using QuantSeq 3’ mRNA Seq Library Prep Kit FWD with Unique Dual Indices for Illumina (Lexogen, cat. no 115.384). Sequencing was performed on an Illumina NovaSeq 6000 platform with NovaSeq 6000 SP Reagent Kit v1.5 100 cycles (Illumina, cat. no. 20028401). RNA seq experiments were conducted with three independent biological replicates.

### RNA seq data analysis

Upon quality trimming using bbduk^45^ (k=13 ktrim=r useshortkmers=t mink=5 qtrim=t trimq=10 minlength=20), fastq files were aligned with STAR^46^ v2.7.3a to the *mm10* mouse reference genome. BAM files were down sampled to 8 million reads with samtools^47^ v1.13 and counted by HTSeq^48^ v2.0.1 (-m union -s no -t exon). Differential expression analysis was performed using DESeq2 ^49^ v1.38.3. Expression of marker genes during RA-driven differentiation were extracted from DESeq2 (counts(dds, normalization = TRUE)) and plotted with Prism v9.2.0. PCA with mean log transformed DESeq2 (rlogTransformation(dds, blind = FALSE)) was plotted with the ggplot2 v3.4.2. PCA mean top 10000 most variable genes was plotted with the ggplot2 v3.4.2. Variant stabilisation transformed counts were plotted for the top 50 most differentially expressed genes with the pheatmap v1.0.12 package.

### Molecular dynamics

All-atom simulations were performed with the GROMACS 2021 MD package^50^ using the EZH2 SET domain (residues 490-751 of EZH2 isoform 2) and the nucleosome from PDB 6WKR as a starting model. Simulations encompassed unmethylated or trimethylated lysines in positions 510, 514 and 515 of the SET domain, either in the presence or absence of the nucleosome, in 100 mM KCl, with two replicates for each case. The parameters for the modified lysines were taken from ^51^*, while Amber forcefields*^52,53^ were used for protein, DNA, and ions. For water molecules, the TIP3P model^54^ was used. Long-range electrostatics were evaluated with particle-mesh Ewald summation,^55^ and all hydrogen bonds were constrained with the LINCS algorithm.^56^ A leap-frog integrator was considered with a 2 fs timestep, and a 1.2 nm cutoff was used for both the electrostatic and Van der Waals interactions. All simulations underwent an initial energy minimization with the steepest descent method,^57^ followed by a 50 ns NpT equilibration with a Parrinello-Rahman barostat^58^ at 1 atm and Nose-Hoover thermostat^59^ at 300 K. Position restraints were applied to the phosphorus atoms of the nucleosomal DNA as well as to the first and last five amino acids of the EZH2 SET domain. 1000 ns simulations were performed for all cases with the position restraints in place. The number of contacts between bridge-helix (EZH2 residues 501- 617) and the H3 histone tail were calculated using GROMACS inbuilt routines. For the helicity analysis, the change % in helicity for the bridge-helix was calculated using GROMACS at an interval of 50 ns for all the cases.

## Supplementary Information

**Supplementary Table 1.**
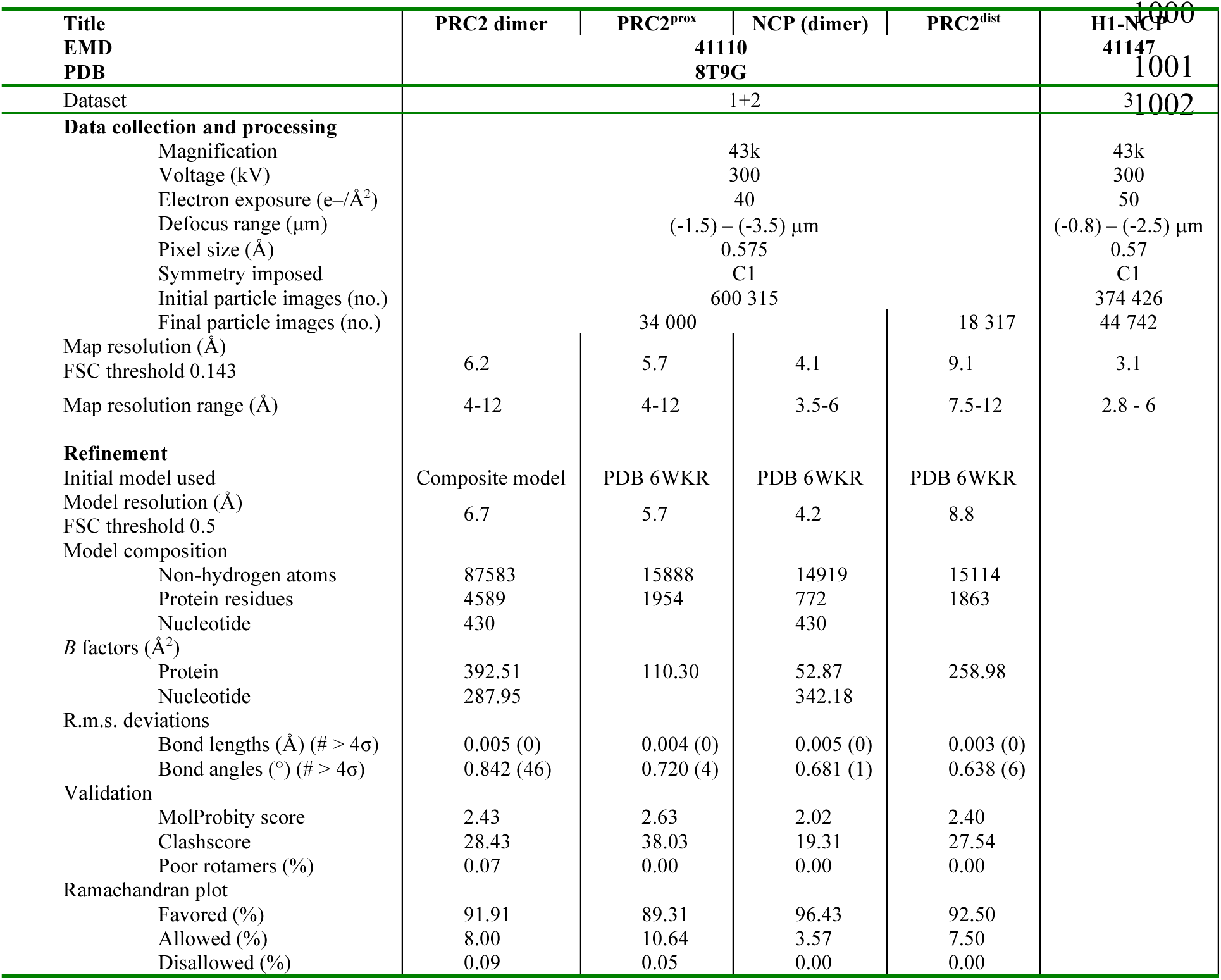
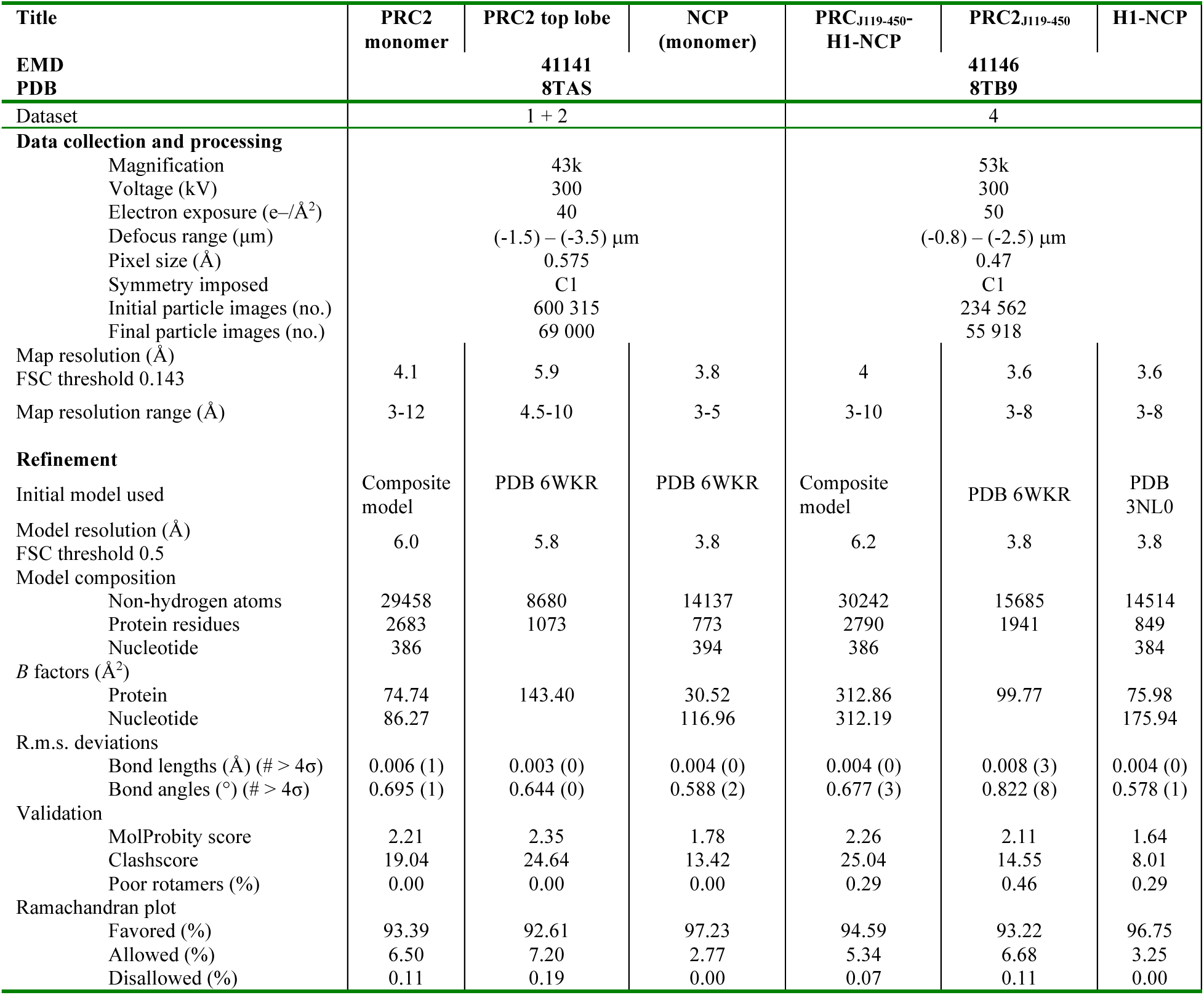
cryo-EM data collection and processing.

**Supplementary Figure 1:**
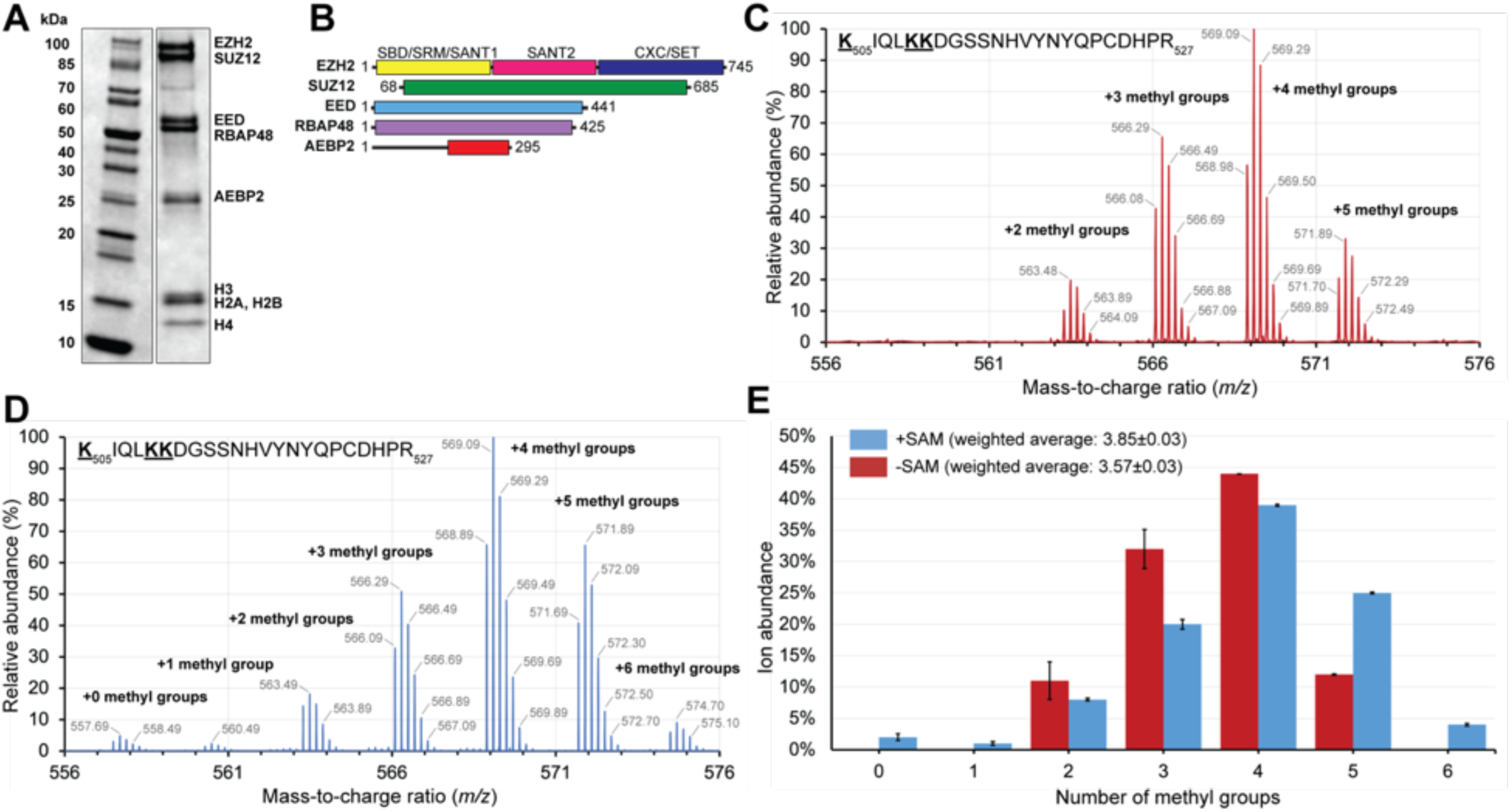
PRC2 purification and automethylation. (A) SDS-PAGE showing subunits of PRC2 and nucleosome used in this study. (B) PRC2 subunits and constructs used in this study. The colors of the domains and subunits are the same as in Fig. 1. (C) Mass spectrometric analysis of automethylated PRC2. Representative high-resolution mass spectra showing detail for the [M+5H]^5+^ ion group for the indicated EZH2 peptide bearing automethylated lysine residues. Cysteines have been alkylated during experiment. (D), like C but after addition of 5mM SAM during protein purification. E, Ion abundance plot showing the number of methyl groups present on average in -SAM and +SAM sample on the automethylation peptide. Error bars show standard deviation (N=2).

**Supplementary Figure 2:**
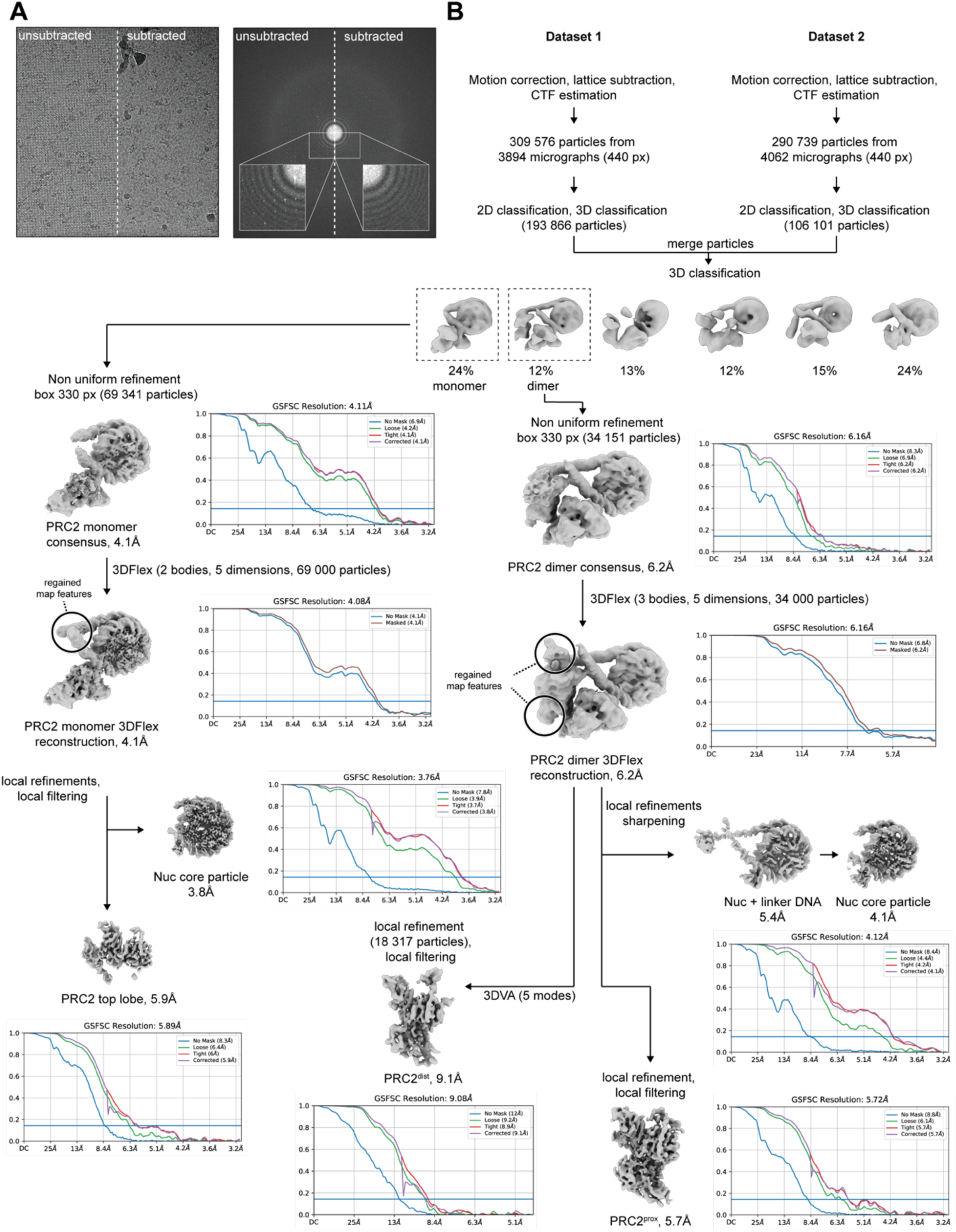
cryo-EM data processing. (A) representative cryo-EM micrograph before and after subtraction of the streptavidin lattice (left) and corresponding Fourier transform showing presence/absence of streptavidin diffraction peaks (right). (B) data processing scheme for PRC2 dimer and PRC2 monomer. See Methods for description.

**Supplementary Figure 3:**
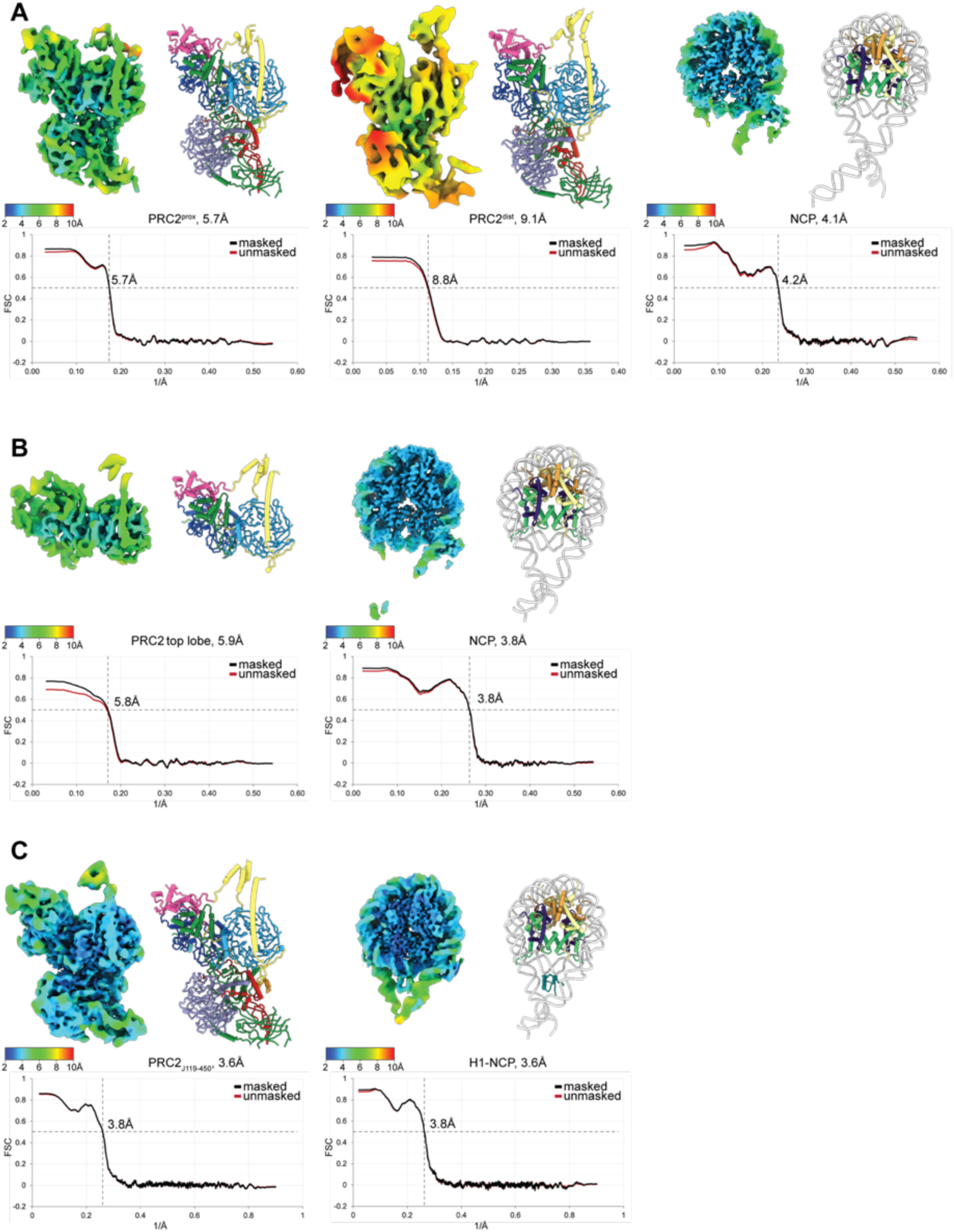
Local resolution and model validation. Local resolution variation and cartoon representation of the final refined model and masked/unmasked map-to-model Fourier shell correlation for (A) PRC2 dimer, (B) PRC2 monomer, (C) H1-nucleosome- PRC2_J119-450_. For each model, the resolution of the cryo-EM model at FSC = 0.143 is provided, while the map-to-model FSC plots show the masked resolution at FSC = 0.5.

**Supplementary Figure 4:**
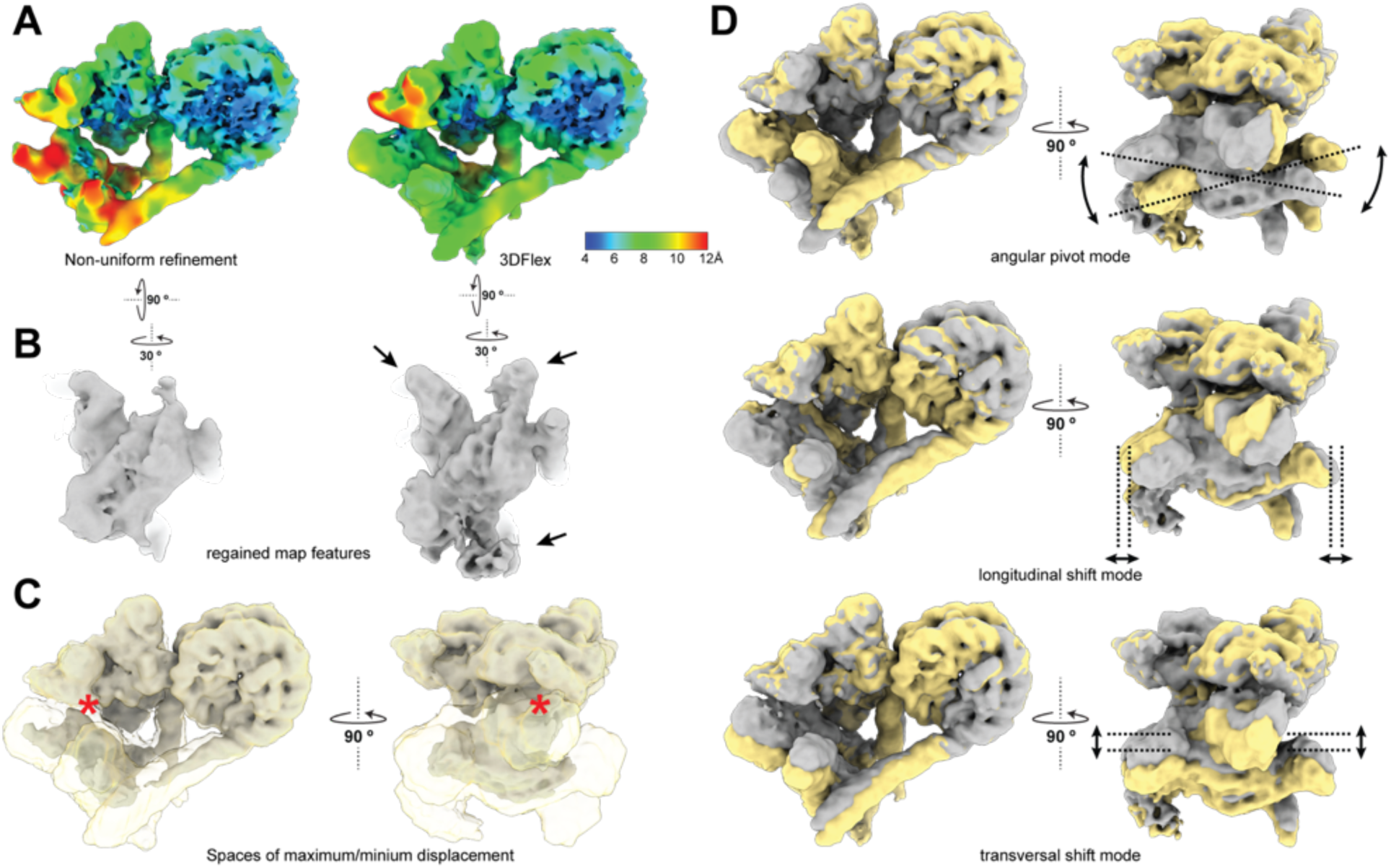
PRC2^dist^ displays large amount of continuous flexibility. (A) Local resolution before (left) and after (right) 3DFlex analysis, showing gain in resolution at the flexible distal PRC2. (B) Comparison of map features at comparable threshold of PRC2^dist^ before (left) and after (right) 3DFlex analysis. Arrows point to parts of the map where the quality has visibly improved. (C) Visualization of continuous flexibility of PRC2 dimer. Yellow, transparent map shows the extent of space sampled by the complex as calculated with 3DFlex.^19^ Grey, solid map shows the region of least variability where density is always present. Red asterisk marks the position of the automethylation loop within the less variable region. (D) Three example modes of movement of the PRC2 dimer, showing pivoting and shifting motions of the distal PRC2.

**Supplementary Figure 5:**
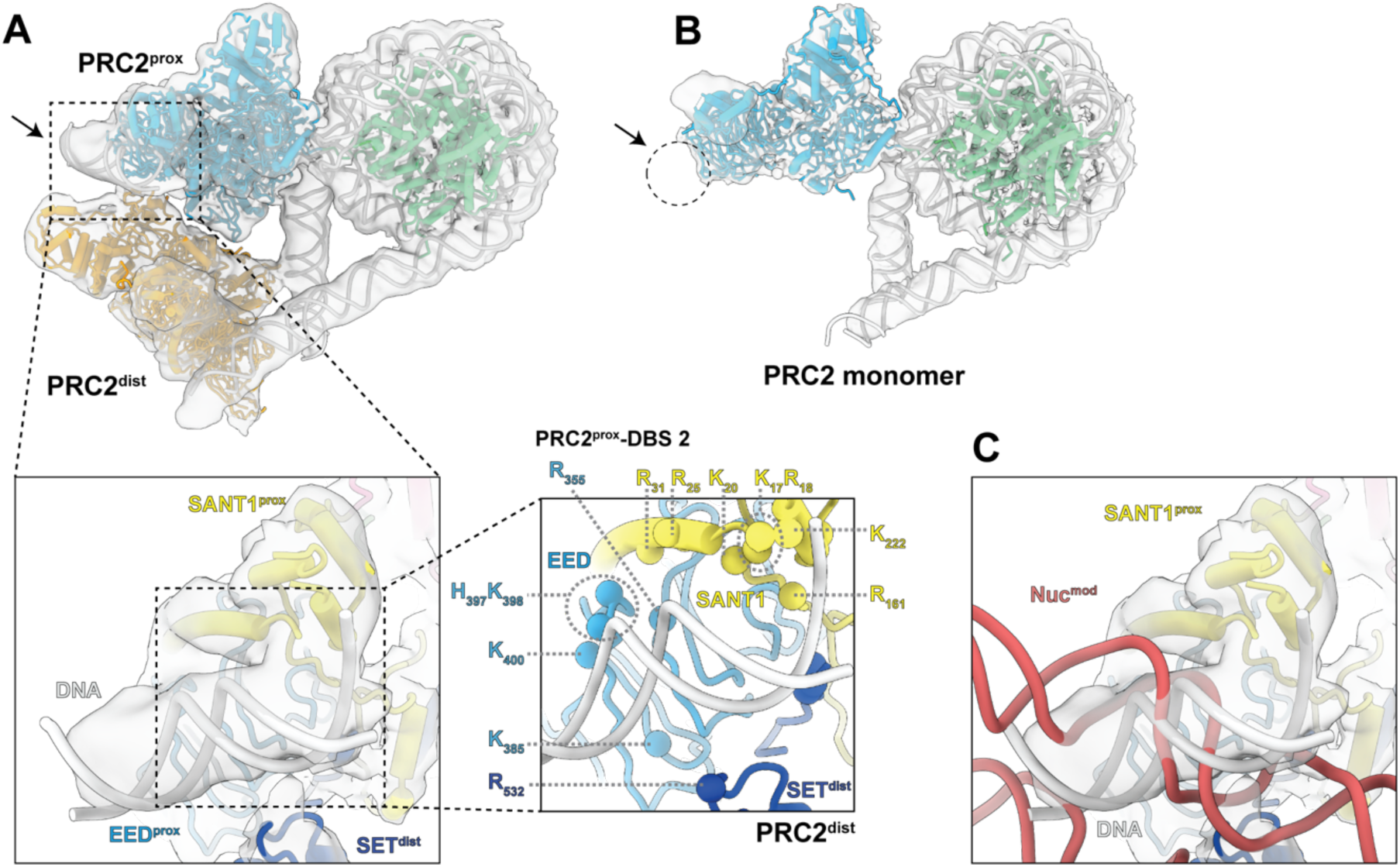
Structural features of the PRC2 dimer. (A) Additional density in the consensus map is indicated with arrows and suggested to be a short piece of dsDNA bound to DBS2 of PRC2^prox^. Below: closeup of extra density with docked DNA model shows how positively charged residues on PRC2^prox^ and PRC2^dist^ could mediate binding. The binding surface would include portions of the PRC2^prox^ SANT1 domain which would only be positioned correctly when PRC2 is active, and the SBD helix is bent. (B) Comparison of (A) with the PRC2 monomer showing the absence of any additional density (arrow and circle). In the PRC2 monomer, the SBD helix is straight (Fig.4C) and the SANT1 domain is not available to mediate interaction with the additional density (C) DNA density and position with respect to an allosteric nucleosome (Nuc_mod_) ^8^ showing that in the presence of an allosteric nucleosome the same DNA binding surface is used (DBS2).

**Supplementary Figure 6:**
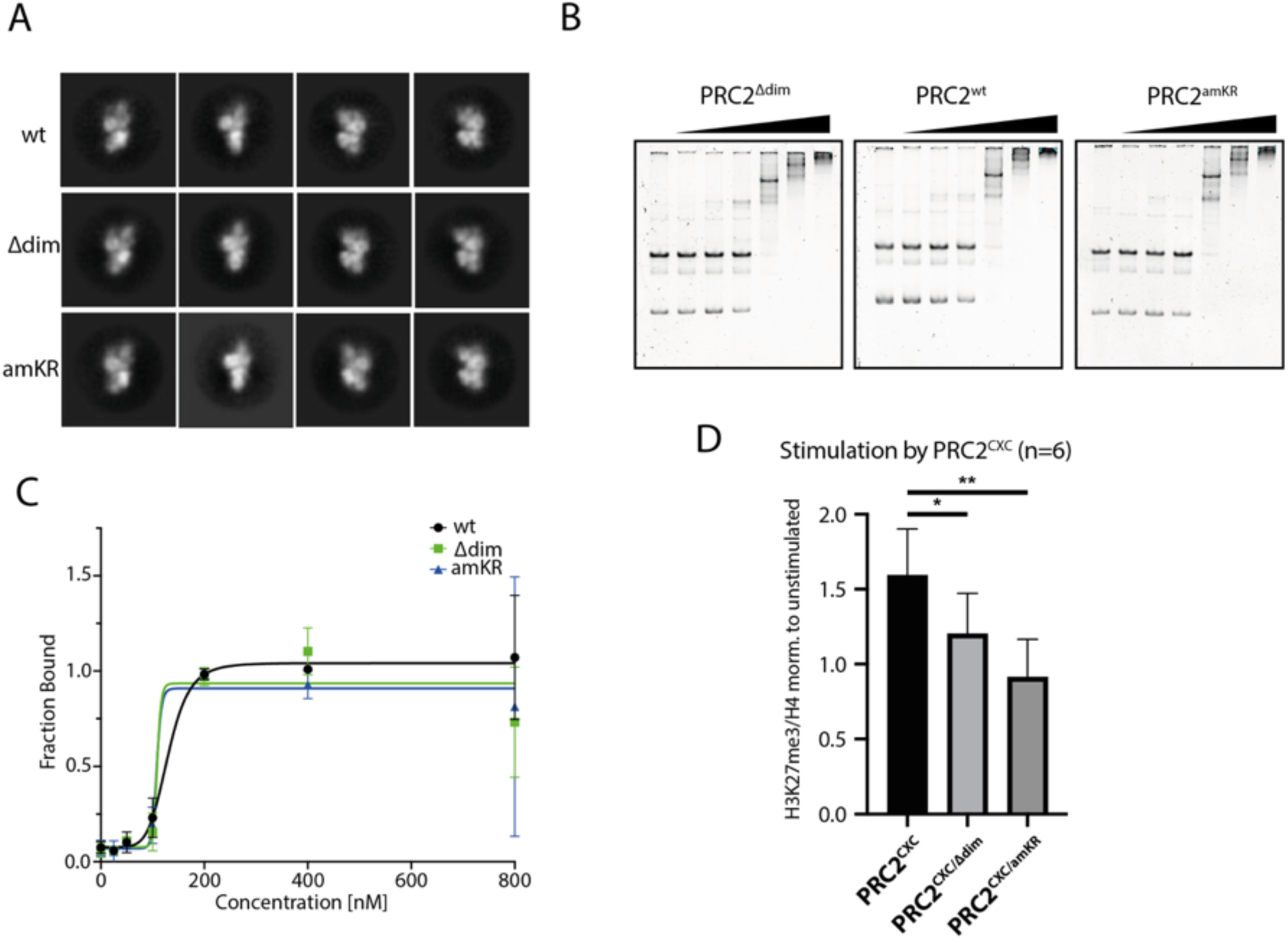
Structure and activity of mutant PRC2 complexes. (A) Representative 2D class averages of WT PRC, PRC2^Δdim^ and PRC2^amKR^ obtained by negative-stain EM. Each row corresponds to the same view for all three complexes. (B) Nucleosome binding of WT PRC2, dimerization and automethylation mutants observed by EMSA. 50 nM mononucleosomes and two-fold titrations ranging from 25–800 nM PRC2. (C) Densitometric quantification of n = 3-6 experiments as shown in B, based on the intensity of free nucleosomes. (D) Quantitative analysis of HMTase activity assays as shown in Fig. 3, based on the band intensities of the H3K27me3 specific antibody relative to the H4 total histone loading controls. H3K27me3 intensities are shown normalized to the activity of unstimulated PRC2^amKR^, i.e. in the absence of PRC2^CXC^.

**Supplementary Figure 7:**
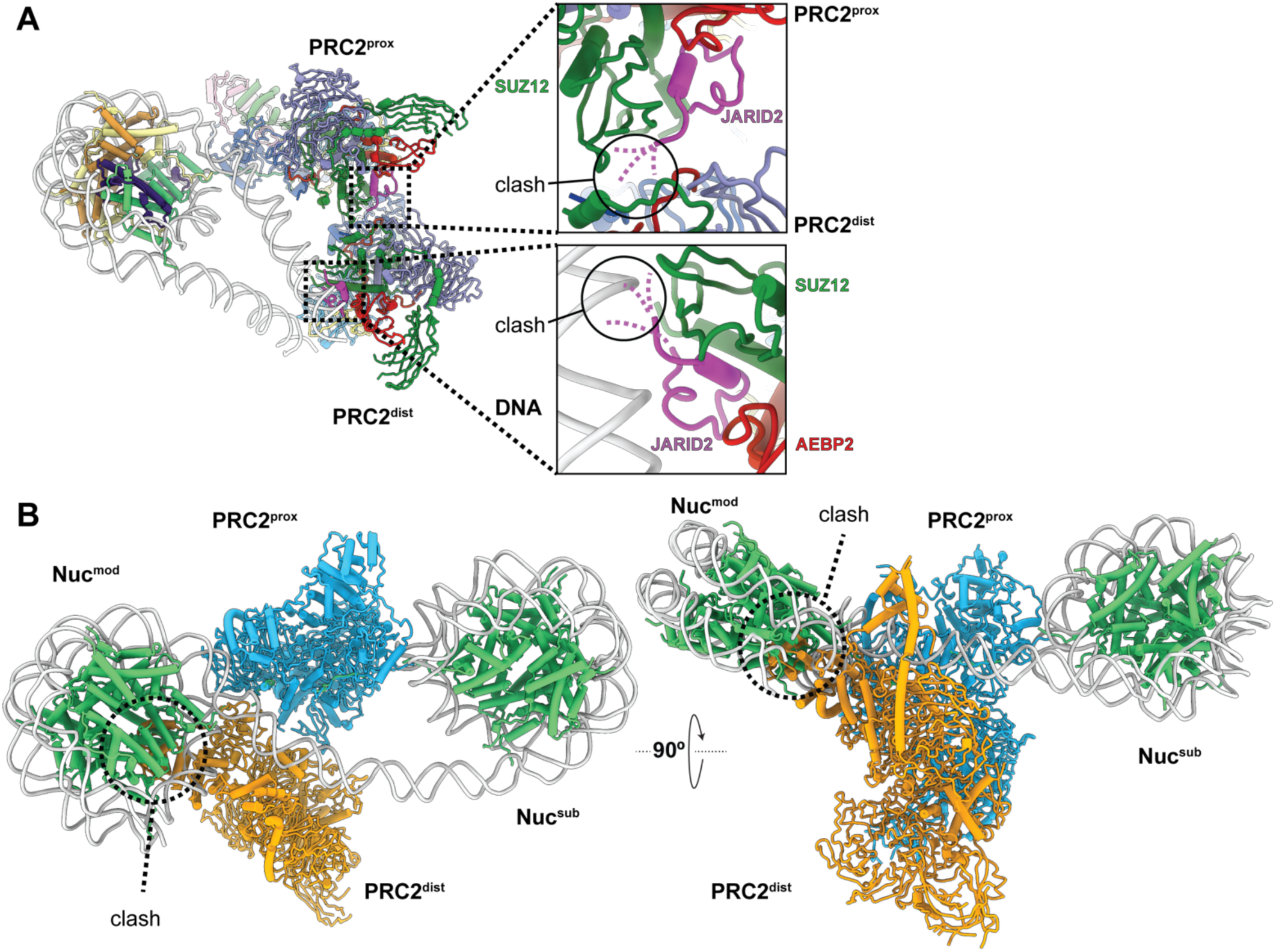
Predicted steric incompatibility of the PRC2 dimer with JARID2 and the dinucleosome. (A) predicted incompatibility between JARID2 and the allosteric PRC2 dimer. The position of JARID2 was derived through alignment of each PRC2 complex with either PDB 6C23 or 6C24. Unmodeled JARID2 segments will likely block the SUZ12-SUZ12 dimer interface (top) and/or prevent SUZ12-DNA binding at DBS3 (bottom). (B) predicted clash between the allosteric PRC2 dimer and the PRC2-dinucleosome. The modified, allosterically activating nucleosome occupies the same space as the allosterically activating PRC2^dist^.

**Supplementary Figure 8:**
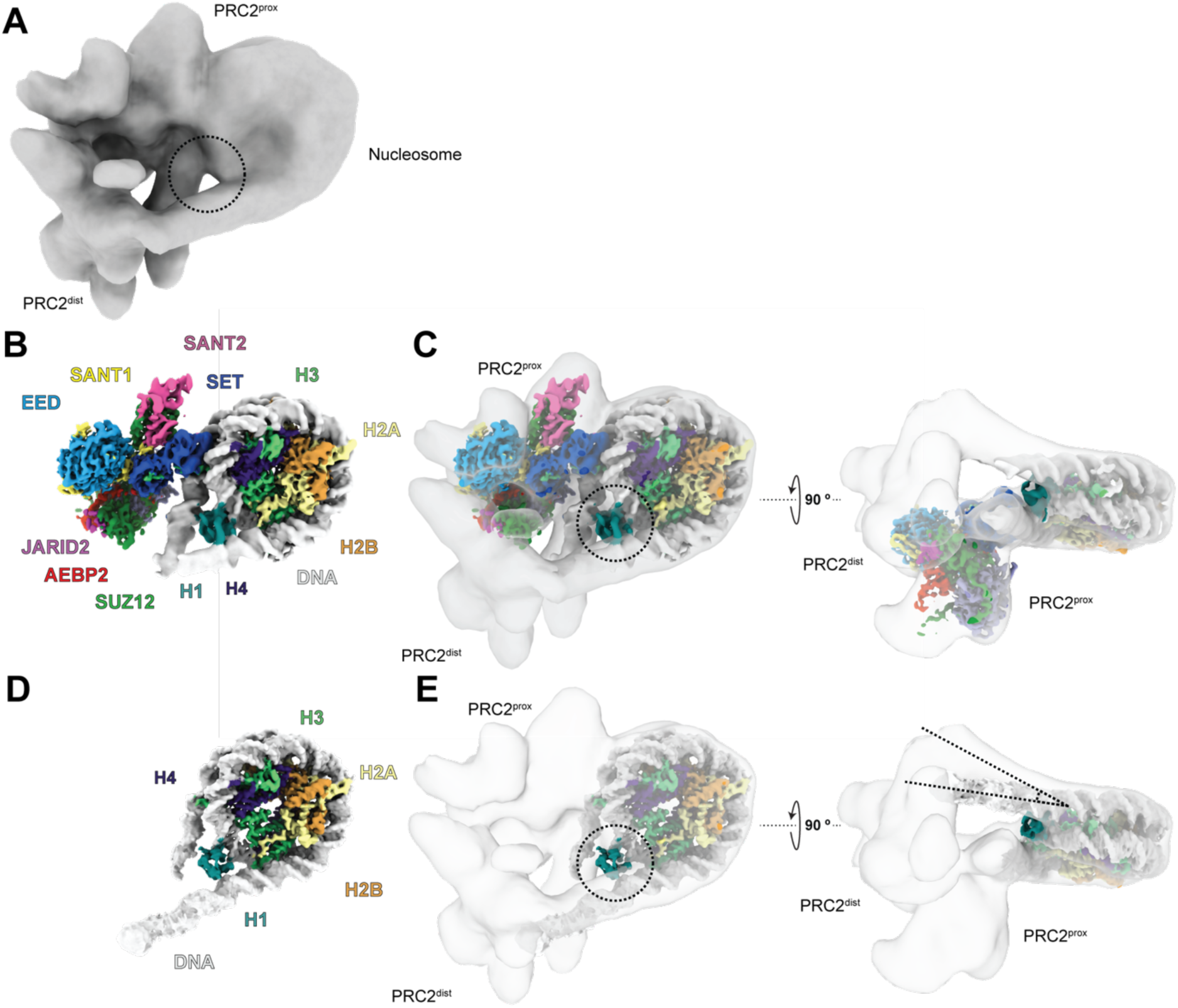
The allosteric PRC2 dimer is incompatible with H1 binding. (A) Cryo-EM reconstruction of PRC2 dimer obtained in the presence of H1 in the sample. The circle indicates where H1 density would have been expected. (B) Cryo-EM reconstruction of PRC2 containing cofactor JARID2 bound to a nucleosome containing H1 (chromatosome) at 3.6 Å resolution. H1 (teal) is bound at the nucleosomal dyad, contacting the linker DNA. (C) Superposition of (A) and (B) shown in two orthogonal views. The circle shows the absence of H1 density in the allosteric dimer. (D) Cryo-EM reconstruction of the chromatosome in the absence of PRC2. (E) Superposition of (A) and (D) based on alignment of H1 in (D) and (C) shown in two orthogonal views. The right panel shows how the trajectory of the chromatosomal linker DNA is incompatible with the DNA geometry seen in the allosteric PRC2 dimer structure.

**Supplementary Figure 9:**
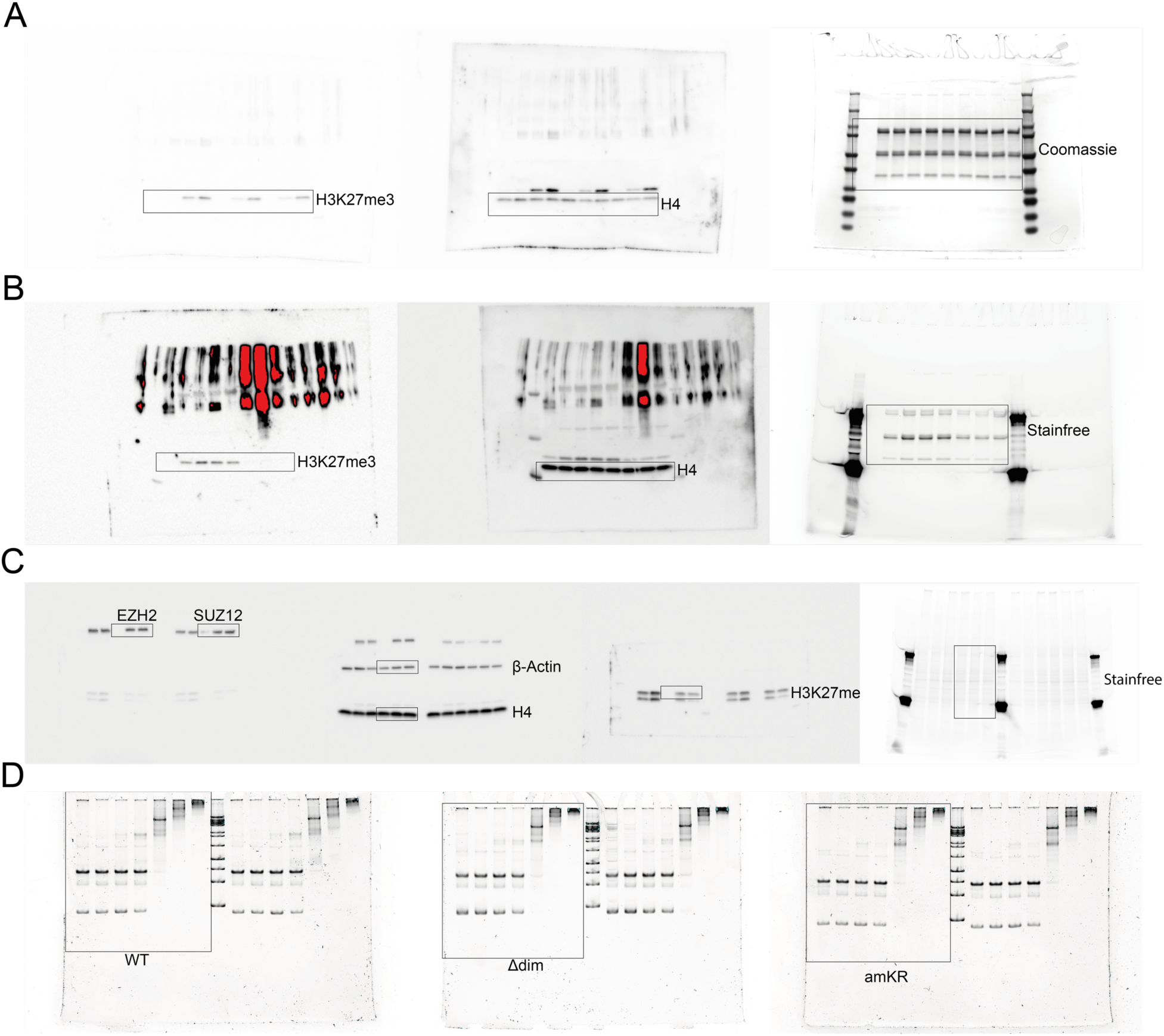
Raw data for Western Blots and native gels. Uncropped images used (A) in Fig. 4C, (B) in Fig. 4D, (C) Fig. 6A and (D) Fig. S6B.

## Notes

### Competing Interest Statement

The authors have declared no competing interest.

